# Temperature dependence of pollen germination in Douglas-fir (*Pseudotsuga menziesii*): A machine learning-based detection of pollen viability from microscopic images

**DOI:** 10.64898/2026.09.10.750703

**Authors:** Hsin-Wu Hsu, Sabrina Zerrade, Soo-Hyung Kim

## Abstract

- *Background and Aims:* Pollen germination and tube growth are critical stages of plant reproduction that are highly sensitive to temperature but remain labor-intensive to quantify. This study aimed to develop a deep learning–based approach for pollen phenotyping and to characterize the temperature dependence of pollen germination and elongation in Douglas-fir (*Pseudotsuga menziesii*).
- *Methods:* A convolutional neural network (CNN) was trained to segment pollen grains and classify germination status from microscopic images. Germination percentage and pollen length were quantified across a range of incubation temperatures from 5 to 40°C. Gamma functions were fitted to temperature response curves to estimate the optimal temperatures for pollen germination and elongation among Douglas-fir populations collected across an elevational gradient.
- *Key Results:* The CNN achieved high segmentation accuracy (intersection over union = 0.846), accurately distinguishing pollen grains from the background but showing moderate accuracy in separating germinated from ungerminated pollen during early elongation. Both pollen germination and elongation exhibited bell-shaped temperature response curves with distinct thermal optima. Pollen elongation consistently reached its optimum at higher temperatures than pollen germination. Estimated optimal temperatures fell within a narrow range, approximately 19 to 23°C. No significant relationship was detected between elevation and thermal optima, although some higher-elevation populations exhibited lower optimal temperatures. Comparisons with previous analyses of three western North American conifers showed that each species occupied a distinct reproductive thermal niche corresponding to the spring temperatures of its native habitat.
- *Conclusions:* Deep learning provides an efficient approach for high-throughput quantification of pollen germination and elongation from microscopic images. The narrow thermal range for reproductive performance suggests that Douglas-fir pollen is sensitive to temperature variation and that warming climates may alter reproductive success. These findings improve our understanding of the thermal sensitivity of conifer reproduction and provide a scalable framework for assessing impacts of climate warming on forest regeneration.

## 1. INTRODUCTION

Pollen viability is a key determinant of reproductive success in conifers, as germination and tube growth represent essential stages leading to fertilization (Steinacher & Wagner, 2012; Taylor & Hepler, 1997). These processes are among the most temperature-sensitive stages of plant reproduction, directly influencing seed production and shaping population persistence under climate change (Hedhly et al., 2009; Hsu & Kim, 2025; Zinn et al., 2010). Traditional pollen viability assessments rely on visual inspection of germinated versus non-germinated grains, a method that is time-consuming, labor-intensive, subjective, and requires extensive expertise (Arias et al., 2017; Battiato et al., 2021; France et al., 2000). While automated pollen grain recognition has been applied in palynology for decades, these applications have largely focused on species or taxonomic classification (Arias et al., 2017; Battiato et al., 2021; France et al., 2000; Gonçalves et al., 2016; Khanzhina et al., 2018; Punyasena et al., 2012; Rostami et al., 2023, 2025). In contrast, methods capable of automatically distinguishing germinated from non-germinated pollen based on morphological traits such as initial elongation remain rare. Therefore, developing machine learning (ML) methods for pollen viability assessment is crucial to improve throughput, accuracy, and reproducibility in pollen phenotyping.

Advances in machine learning have significantly enhanced pollen image analysis by combining feature extraction with classification algorithms (Arias et al., 2017; Battiato et al., 2021; Gonçalves et al., 2016; Rostami et al., 2023, 2025). Early approaches relied on geometric, textural, and morphological descriptors, including Bag of Visual Words (BOW) and Color, Shape, and Texture (CST) features (Arias et al., 2017; Gonçalves et al., 2016). Classification was then performed using algorithms such as Support Vector Machines (SVMs), decision trees, and K-nearest neighbors (KNN), often achieving accuracies above 80% (Arias et al., 2017; Gonçalves et al., 2016). More recently, deep learning approaches, particularly convolutional neural networks (CNNs), have transformed pollen analysis by automatically extracting features from raw images and achieving accuracies exceeding 95% (Battiato et al., 2021; Khanzhina et al., 2018; Rostami et al., 2023, 2025). These advancements are increasingly supported by high-performance machine learning libraries such as PyTorch, which offer the dynamic execution needed for flexible model development (Paszke et al., 2019). Architectural innovations have also played a critical role; for example, the inception architecture enables deeper networks through efficient multi-scale processing and dimensionality reduction (Szegedy et al., 2015). Furthermore, the integration of attention gates (AGs) in models like the Attention U-Net allows for more precise segmentation by suppressing irrelevant background data and focusing on salient biological structures (Oktay et al., 2018). Current research highlights the efficacy of transfer learning, e.g., adapting pre-trained models such as ResNet101 and ResNeSt-110 to pollen datasets, which deliver faster results and higher accuracy than traditional models built from scratch (Rostami et al., 2023, 2025). These advanced models have proven capable of distinguishing between morphologically similar conifer genera, such as *Abies*, *Picea*, and *Pinus*, achieving test accuracies as high as 99% (Rostami et al., 2025). Although CNNs have been applied successfully to genus and species-level pollen recognition, their use in identifying germination status remains limited, representing an important methodological frontier for pollen biology.

Douglas-fir (*Pseudotsuga menziesii*) pollen germination presents an ideal case study for applying ML-based image recognition methods. Unlike many other gymnosperms and angiosperms, Douglas-fir pollen does not develop a conventional tube in the initial phase but instead elongates as the exine ruptures and the pollen cytoplasm emerges (Ho, 1968; Owens et al., 1981). The process of Douglas-fir pollen germination takes about 2 weeks and involves two phases: during the initial phase, the pollen swells by taking up moisture, then elongates and expands, and the second phase, the production of the pollen tube, requires additional factors (Dumont-BéBoux & Von Aderkas, 1997). In addition, Douglas-fir’s exceedingly slow *in vivo* growth requires about nine weeks from pollination to fertilization (Allen, 1943; Ho, 1968). *In vitro*, pollen germination is typically defined as the tube elongation exceeding the pollen grain diameter, a criterion that is morphologically distinct and readily quantifiable (Shivanna & Rangaswamy, 1992; Varis et al., 2010). However, Douglas-fir pollen is difficult to classify manually because of its irregular morphology, delayed development (no initial tube growth), and slow growth *in vivo* (Webber & Bonnet-Masimbert, 1993; Webber & Painter, 1996). Despite the difficulties, the transition of Douglas-fir pollen from a dormant grain to an elongated grain provides measurable geometric features (e.g., shape, contour, size ratio) that machine learning methods are highly effective at extracting and quantifying. Automated classification provides a scalable and objective approach to detect initial elongation, bypassing the laborious effort and subjectivity of manual counting. Thus, Douglas-fir pollen morphology and growth dynamics provide an effective test system for advancing ML-based phenotyping of conifer reproductive traits.

Temperature is the major environmental factor influencing conifer reproduction, particularly during the highly vulnerable phases of pollen germination and tube growth (Hsu & Kim, 2025). Experimental studies demonstrate that pollen viability is strongly shaped by hydration conditions, storage environment, and incubation temperature (Hsu & Kim, 2025; Siregar & Sweet, 2000), with optimal germination typically occurring in environments closely related to their natural habitats (Hsu & Kim, 2025). Extremely high or low temperatures reduce germination percentage and elongation, which means constraining seed set and fertility (Moody & Jett, 1990; Webber et al., 2005). These physiological responses suggested that Douglas-fir reproductive timing and success are also directly linked to temperature-dependent phenology, pollen competition, and seed production. Furthermore, variation in the thermal requirements of pollen germination influences conifer species distributions, with climate change expected to reduce pollen performance, alter reproductive synchrony, and shift population dynamics (Bykova et al., 2012; Hsu & Kim, 2025; Morin et al., 2007). Therefore, understanding the temperature dependence of Douglas-fir pollen germination is essential for predicting both reproductive outcomes in managed seed orchards and population resilience under future climate scenarios.

This work builds upon the previous study, Hsu & Kim (2025), which demonstrated the temperature dependence of pollen germination and tube growth in conifer species. The foundation laid by this previous work emphasizes the importance of characterizing temperature-induced reproductive failure in conifers. The overall goal of the current study is twofold: (1) to develop a robust, automated ML platform capable of rapidly and accurately identifying Douglas-fir pollen germination based on initial elongation, thus significantly improving the throughput and objectivity of viability assessment, and (2) to leverage this automated method to precisely quantify the temperature dependence of Douglas-fir pollen germination and pollen elongation as a way of assessing climate change impact on reproductive success. Our primary hypothesis is that a CNN model trained on a large dataset of labeled Douglas-fir pollen images will be an effective means to determine pollen viability and growth with high accuracy based on initial elongation during the first stage, providing precise and reproducible measures of germination percentage and pollen length that substantially reduce the time and labor required for manual counting. Furthermore, we hypothesize that automated measurements will allow us to characterize the temperature dependence of Douglas-fir pollen and compare with our previous findings in other conifers co-occurring in the Pacific Northwest region (Hsu & Kim, 2025), providing critical insights into reproductive ecology under climate change.

## 2. MATERIALS AND METHODS

### 2.1 Pollen collection sites

We collected pollen of Douglas-fir (*Pseudotsuga menziesii* (Mirb.) Franco; PSME) at Tyee Mountain (350–2000 m; Entiat) and Slate Peak (1500–2200 m; Mazama) in Washington State, USA (Figure 1). Nine pendant temperature/light data loggers (UA-002-64, Onset HOBO, Bourne, MA, USA) were deployed to record hourly mean air temperatures at Tyee Mountain along the elevational gradient and seven loggers at Slate Peak along the elevational gradient from October 2020 to June 2021 (Figure 1). The recorded spring mean temperature (April, May, and June) range at the pollen collection sites for Douglas-fir is 10.3–12.6 °C. The monthly mean temperatures along the elevational gradients in Tyee Mountain and Slate Peak were shown in Hsu & Kim (2025). Douglas-fir naturally occupies a wide range of climatic conditions and elevations from sea level to 1520 m in Washington State, USA, extending from 19 to 55° N in North America, and pollen is shed during March and April in the warmer part of the range and as late as May or early June in the colder areas (Burns & Honkala, 1990). The pollen maturity date of Douglas-fir is from late April to May at our sites. We collected Douglas-fir mature male cones before the scales opened and shed pollen from May 1 to May 23, 2021, at 30 elevations from 800 to 1500m. The pollen was collected from multiple cones in the same tree at each elevation. After the pollen collection, we placed male cones at room temperature to dry for 5 days. We then collected the pollen grains and moved them to the refrigerator (4 °C) before the viability tests.

**Figure 1.**
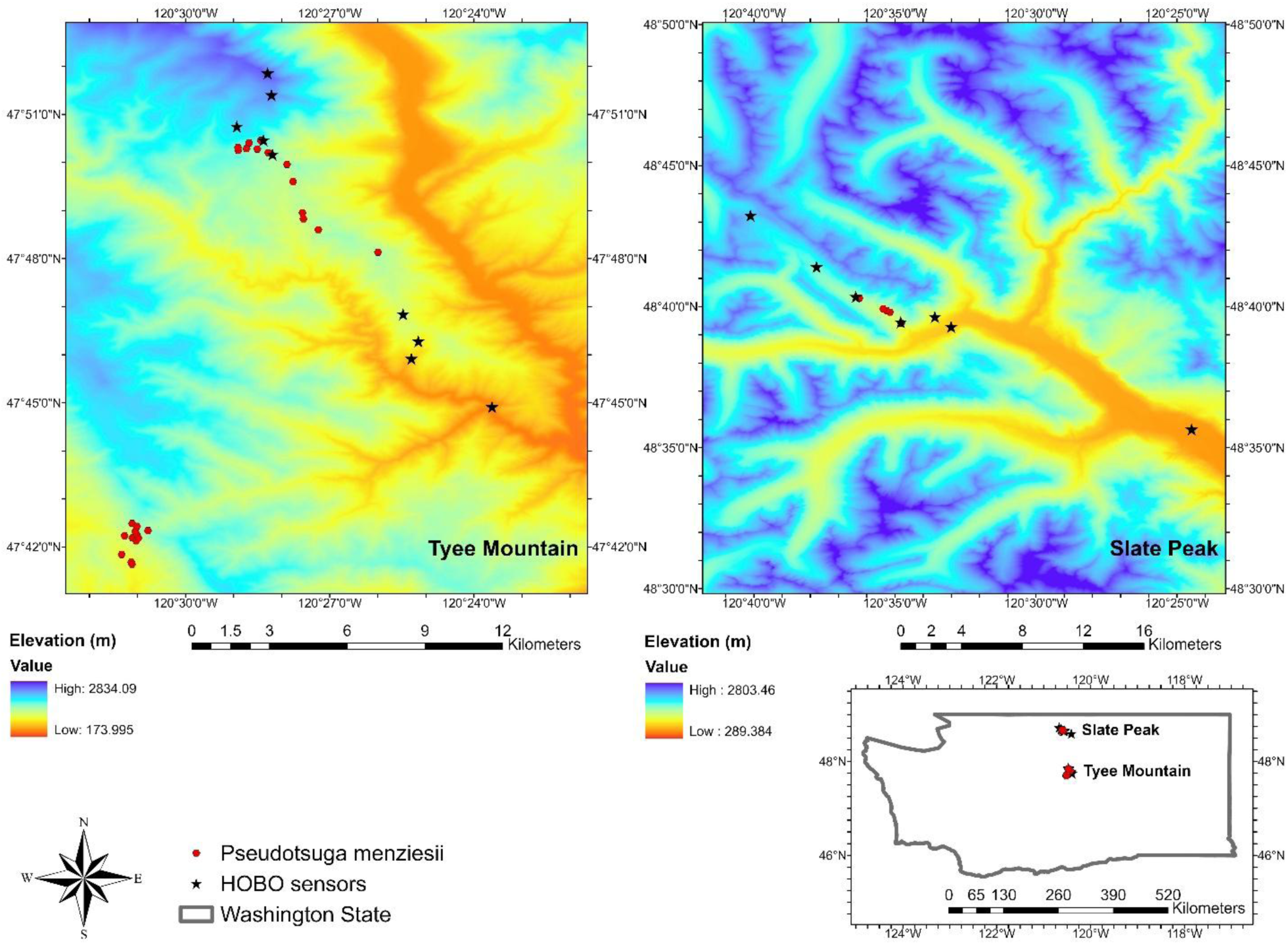
Temperature data and pollen of *Pseudotsuga menziesii* were collected from the elevational gradients at Tyee Mountain and Slate Peak in Washington State, USA.

### 2.2 Pollen germination test

Pollen germination tests were conducted from July to September 2021. A suspension culture method (Shivanna & Rangaswamy, 1992) was used to test pollen germination. We first cultured pollen with the germination medium of Brewbaker & Kwack (1963), which was autoclaved and supplemented with thiamine, riboflavin, and ascorbic acid (Hsu & Kim, 2025; Varis et al., 2011). This recipe works well in Scots pine and is close to what Tushabe & Rosbakh (2021) suggested for the gymnosperms. The vials were also autoclaved before the test, and 5 ml of culture media and 25 mg of pollen grains from each elevation were added to each vial. They were then placed on CO-Z orbital shakers (80 revolutions per minute; Amazon, Seattle, USA) in G-1000 growth chambers (Conviron, Winnipeg, Canada) with temperatures from 5 to 40°C with an interval of 5°C for four days without light (Hsu & Kim, 2025; Parantainen & Pasonen, 2004; Parantainen & Pulkkinen, 2002). The pollen grains of all elevations were placed in the same chamber at each temperature. Three chambers were used in the test. Two of them were set to two temperature treatments for four days. We then reset another two chambers to another two temperature treatments for another four days. We repeatedly reset the temperature treatments until eight temperatures had been completed. Three test rounds (replicates) of each elevation and temperature treatment were conducted in different chambers. At the end of the four days in each round, 3 ml of 0.1% (w/v) aniline blue-lactic acid-glycerol-water solution was added to each vial to arrest pollen growth and stain the pollen blue (Hsu & Kim, 2025; Parantainen & Pasonen, 2004; Parantainen & Pulkkinen, 2002). The vials were then moved to a fridge (4°C), waiting for imaging.

In each test round, two subsamples were collected from each vial and considered to be repeat observations. The six subsamples of each elevation were used in statistical analysis and model fitting. Each subsample was placed on a microslide and covered with a cover glass. We then took images of these samples as soon as the test was completed, under the Trinocular Stereo Microscope SM-2T-LED and 10MP USB 3.0 Color CMOS C-Mount Microscope Camera MU1003 (AmScope, Irvine, California, USA) with a magnification of x14 to x40 to monitor pollen germination. The imaging process was done from July 2021 to October 2021.

### 2.3 Pollen counting

A pollen tube emerges within a few days in *Pinus* and *Picea*. However, in *Pseudotsuga*, the pollen releases itself from its tough outer casing (exine) and is then covered only by a thin inner layer (intine). The pollen tube does not start growing until the pollen has moved close to the central part within the ovule (nucellus), and this may take several weeks after the initial pollination event (Owens et al., 1981; Takaso & Owens, 1994). *Pseudotsuga* exhibits a unique pollen germination pattern characterized by a two-step process. The initial step involves the elongation of pollen, followed by the subsequent formation of pollen tubes. Despite several attempts having been made to induce pollen tube formation in *Pseudotsuga* (Fernando et al., 2005), the process remains poorly understood due to the difficulty in inducing pollen tube growth *in vitro* (Owens & Morris, 1990). Therefore, manual determination of Douglas-fir pollen germination percentage followed a different protocol; the pollen was classified into Classes 1 to 4, and the total number of pollen grains scored in both Classes 1 and 2 determined the overall percent germination (Webber & Bonnet-Masimbert, 1993; Webber & Painter, 1996; Table 1), and Classes 1 and 2 were counted by a CNN-based pollen counter.

**Table 1.** Douglas-fir pollen classification (Webber & Bonnet-Masimbert, 1993; Webber & Painter, 1996).

| Classification | Criteria |
| --- | --- |
| Class 1 | Elongating grain is >2x original diameter. |
| Class 2 | Elongating grain is <2x original diameter. |
| Class 3 | Grain is hydrated (original diameter) but not elongating or showing signs of plasmolysis. |
| Class 4 | Damaged; plasmolysis is evident (includes Class 1, 2, and 3 grains). |

### 2.4. CNN algorithm implementation and development

Semantic segmentation algorithm labels each pixel in an image with a class. Because each pixel is labelled, the boundaries of the objects in the image can be clearly identified and their dimensions determined. This makes the approach useful not just for counting pollen, but also for measuring it. The Attention U-Net is a CNN architecture developed for medical image segmentation (Oktay et al., 2018). The inception network is another CNN architecture formulated to tackle image detection tasks while limiting the computational budget of a deep neural network (Szegedy et al., 2015). A model architecture that combines the features of these two models was developed for stomata counting and measurement (Gibbs et al., 2021). This model architecture was selected as the basis for the model for pollen classification because it was capable of achieving high accuracy in a similar semantic segmentation task.

To train the model, a set of 130 pollen images collected from various elevations and temperatures was manually annotated to create segmentation masks. Each pixel in the image was annotated as one of three categories: the background class, ungerminated pollen (classes 3& 4), and germinated pollen (classes 1 & 2), using the software PixelAnnotationTool (Bréhéret, 2017). The images were divided into training, validation, and testing sets, with approximately 80% of the data for training, 10% for validation, and 10% for testing. As part of the training process, random image augmentations, including cropping, scaling, and filtering, were applied to increase the size and variety of the training set.

A CNN may have millions of weights that are used as constants in the functions making up the model. Training a CNN requires fitting these values to find the best function to transform the original image into a labelled one. To choose the initial weights at the start of the training process, Xavier Normal initialization was used for fast convergence and a consistent training process. The model was trained using the stochastic gradient descent (SGD) optimizer, a method used to minimize loss and find the optimum values for weights. This method involves randomly selecting an example in the training set in each iteration and using it to calculate a gradient that is used to update the values of the weights. SGD is a commonly used optimizer and is expected to approach the minimum loss after many iterations.

Training began with a learning rate of 0.5 for 200 epochs using a momentum of 0.9 and a batch size of 5. After the validation intersection over union (IOU), a measure of the overlap between the predicted masks and the ground truths that is commonly used in segmentation tasks, plateaued at approximately 0.699, the resulting weights were used to initialize subsequent rounds of training with progressively lower learning rates. The learning rate was reduced to 0.1 for approximately 30 epochs, then to 0.05 for 25 epochs, and finally to 0.005 for 30 epochs, resulting in a final validation IOU of 0.846 (Supplementary Data Figure S1). Various combinations of hyperparameters were evaluated during model development, including different learning rates, batch sizes, weight initialization methods, and the Adam optimizer, but SGD provided better performance.

### 2.5 Workflow for image analysis

After the model is trained, it can predict the locations of the pollen grains in the image, creating a labelled segmentation mask. A Python program was developed to measure the length of each individual pollen grain in an image based on this mask and classify it as germinated or not. To determine pollen grain length, contours were identified in the predicted segmentation masks using the OpenCV package for Python and were considered the boundary between the pollen grain and the background. The major axis was measured in pixels and converted to micrometers using the known image scale.

Two methods of determining pollen germination were evaluated and are referred to as length-based and ratio-based methods (Figure 2). The length-based method compared the length of each pollen grain with the average pre-germination length for that elevation. A 95% confidence interval was constructed using the mean length and standard deviation of ungerminated pollen for each elevation. Pollen grains with lengths within this interval were classified as ungerminated, whereas those exceeding the upper bound were classified as germinated. The remaining pollen grains were considered damaged or mismeasured and were excluded from germination percentage calculations. The ratio-based method compared the length-to-width ratio of each pollen grain, based on the assumption that pollen becomes more elongated following germination. The average length-to-width ratio of ungerminated pollen was calculated from seven images of ungerminated pollen, and a 95% confidence interval was constructed. Pollen grains with ratios exceeding the upper confidence limit were classified as germinated.

**Figure 2.**
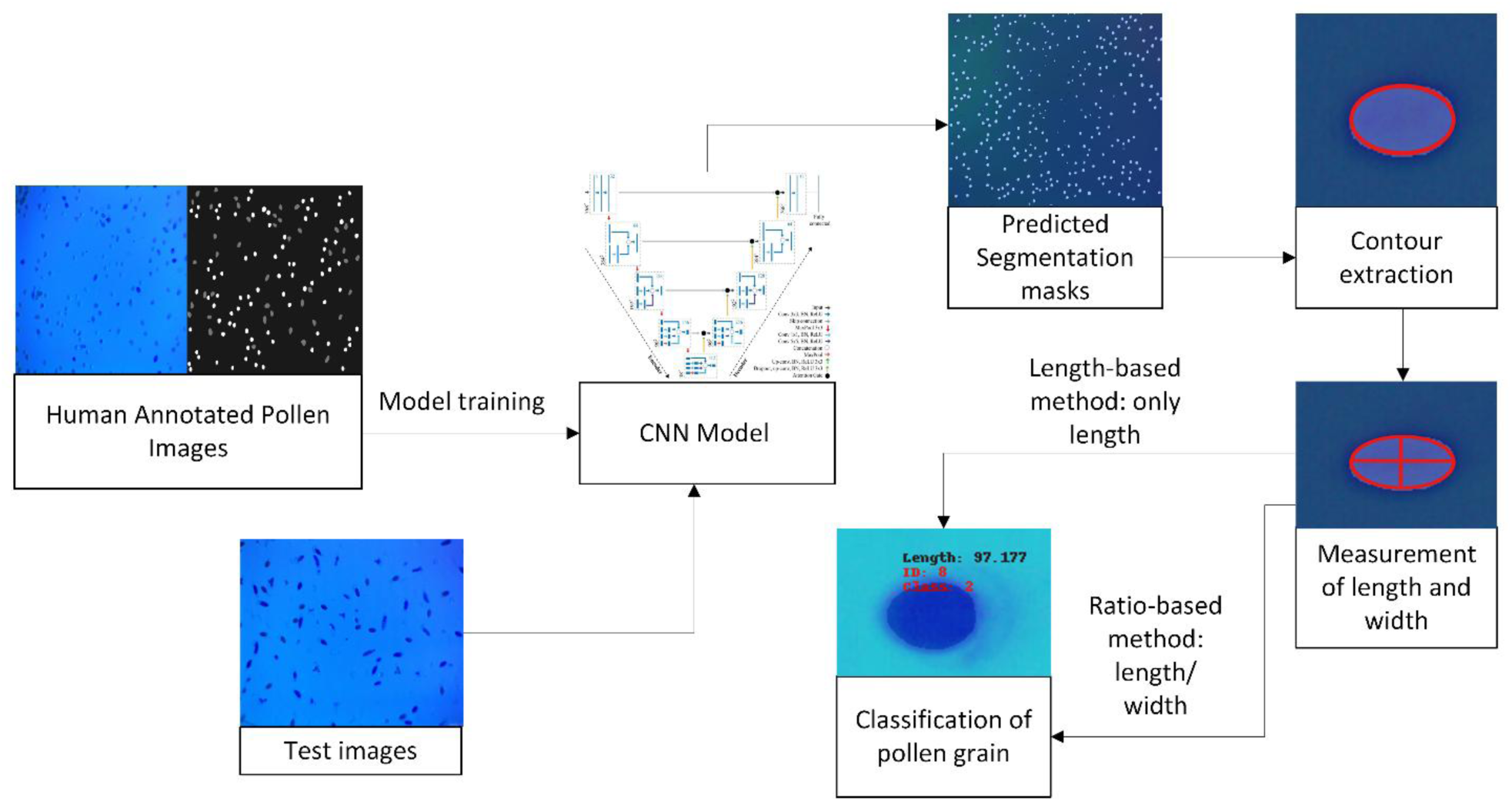
Workflow for analysis of images. First, human-annotated pollen images are used to train the CNN model. The image shows the architecture of the CNN (Gibbs et al., 2021). After training, Gibbs et al.’s model is used to segment the test images, producing segmentation masks with each pixel labelled as germinated, ungerminated, or background. The contours are then extracted from the mask, identifying the boundaries of the pollen. The major and minor axes of the contours are used to determine the length and width of the pollen grain. Then, two possible methods were tried to classify the pollen grains. The length-based method compared the length to a 95% confidence interval based on the mean length before germination to classify the pollen as germinated or not. The ratio-based method compared the ratio between length and width to a 95% confidence interval centered around the mean ratio before germination to classify the pollen.

### 2.6 Modeling pollen germination percentage

We applied the Gamma function (eqn. 1) as used by Hsu & Kim (2025) to evaluate the temperature dependence of pollen germination percentage (GP) and pollen length (PL).

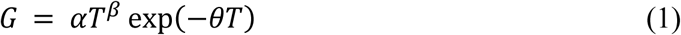

In eqn. (1), *G* is the response variable (pollen germination percentage [unit %], *T* is the explanatory variable (temperature [unit °C]), α (no unit) determines the overall scale or height of the curve, β (no unit) controls the shape of the curve, particularly the steepness of its rise, and θ (no unit) affects the curve’s position along the x-axis (horizontal shift) and influences the curve’s decay rate (Hsu & Kim, 2025). As mentioned in Section 2.3, Douglas-fir’s pollen will not grow tubes until several weeks after the initial pollination. Therefore, the PL here is the length of the elongated pollen, which began from the original pollen diameter. We added an intercept to the Gamma function to fit PL (eqn. 2) to address this difference from Hsu & Kim (2025).

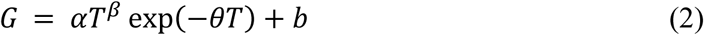

In eqn. (2), the definitions of *G, T,* α, β, and θ are the same as in eqn. 1; *b* is the intercept (pollen original diameter [unit nm]) of the Gamma function.

The Michaelis-Menten equation (eqn. 3) was used to model the GP optimal temperature (*T_opt_GP_*) and PL optimal temperature (*T_opt_PL_*) versus spring mean temperatures across Douglas-fir. We also added three other species in Hsu and Kim (2025) to the figure to show the relative thermal niche of each species.

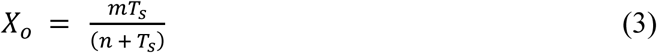

In eqn. (3), *X_o_* is the response variable (optimal temperatures for each species and elevations, unit: °C), *T_s_* is the explanatory variable (spring temperatures at different sites, unit: °C), *m* is the higher asymptote of *X_o_* (*T_s_* → *X_o_*, unit: °C), and *n* is the *T_s_* value giving a response equal to *m*/2 (unit: °C). We restricted the spring temperature to no greater than the optimal temperature (*T_s_* ≤ *X_o_*) in this equation. A summary of each variable and parameter in eqn. (1), eqn. (2), and eqn. (3) is shown in the Appendix.

### 2.7 Statistical analysis

To measure the performance of the model during training and testing, several different metrics were used. When considering two possible outcomes, a correctly predicted positive outcome is called a true positive (TP), while an incorrectly predicted positive outcome is called a false negative (FN). Similarly, a correctly predicted negative outcome is called a true negative (TN), and an incorrectly predicted negative outcome is called a false positive (FP). The proportions of true and false predictions by class can be shown in a figure called a confusion matrix. An accuracy score can be calculated by taking the number of correct predictions divided by the total number of predictions.

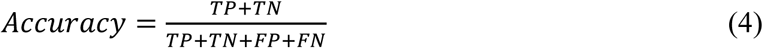

Precision is used to describe what proportion of positive labels are true positives and is calculated using the formula:

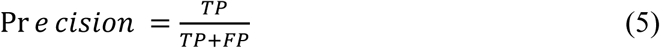

Recall is used to describe what proportion of positive outcomes are correctly labelled and can be calculated as:

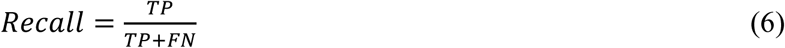

The F1-score depends on both the precision and recall of a model, according to the formula:

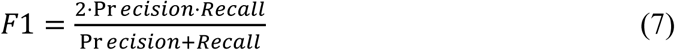

To achieve a high F1-score, a model needs both high precision and recall. To guide training, IOU was used. If comparing two sets X and Y, IOU is defined as:

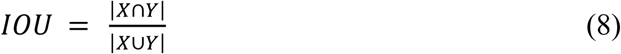

In terms of the elements of the confusion matrix, this can be written as:

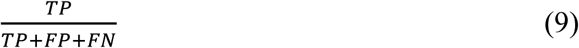

The Python library PyTorch was used to implement and train the model (Paszke et al., 2019). The program to identify and measure pollen for this project was written using the code for the StomataGSMax measurement tool as a base (Gibbs et al., 2021). The code from the stomatal measurement project is available publicly on GitHub at https://github.com/DrJonoG/StomataGSMax. The full code with modifications to make it more applicable for this task is available on https://github.com/uwkimlab/PollenCounter.

We used simple linear regression to fit the *T_opt_GP_* and *T_opt_PL_* to different elevations within the same species. We also used a one-way analysis of variance (ANOVA) and Tukey’s honest significance test (HSD) to test the *T_opt_GP_* and *T_opt_PL_* among Douglas-fir and other species in Hsu and Kim (2025). A t-test was used to compare the differences between Douglas-fir’s *T_opt_GP_* and *T_opt_PL_*, and the “emtrends” function in the “emmeans” package (Lenth, 2022) was used to test the one-to-one relationship between *T_opt_GP_* and *T_opt_PL_*. All analyses were conducted in the R 4.1.2 programming language (R Core Team, 2021).

## 3. RESULTS

### 3.1 CNN model performance for pollen segmentation and gemination detection

The CNN model was used to generate predictions on the test set images, and the IOU for each class was calculated for each image and averaged. The background class had an IOU of 98.74%, the germinated class had 64.33%, and the ungerminated class had 63.14%. The IOU measures suggest that the model has a high accuracy when distinguishing between the background and the pollen grains, but moderate accuracy when distinguishing between the germinated and ungerminated pollen.

For images in the test set, germination percentages measured by humans and the predicted germination percentage are strongly correlated with a close to one-to-one relationship (Figure 3a). The machine learning model tends to underestimate the germination percentage of these test images. This could indicate that the lengths of the individual pollen grains are being underestimated by the model, causing more of them to be counted as non-germinated.

**Figure 3.**
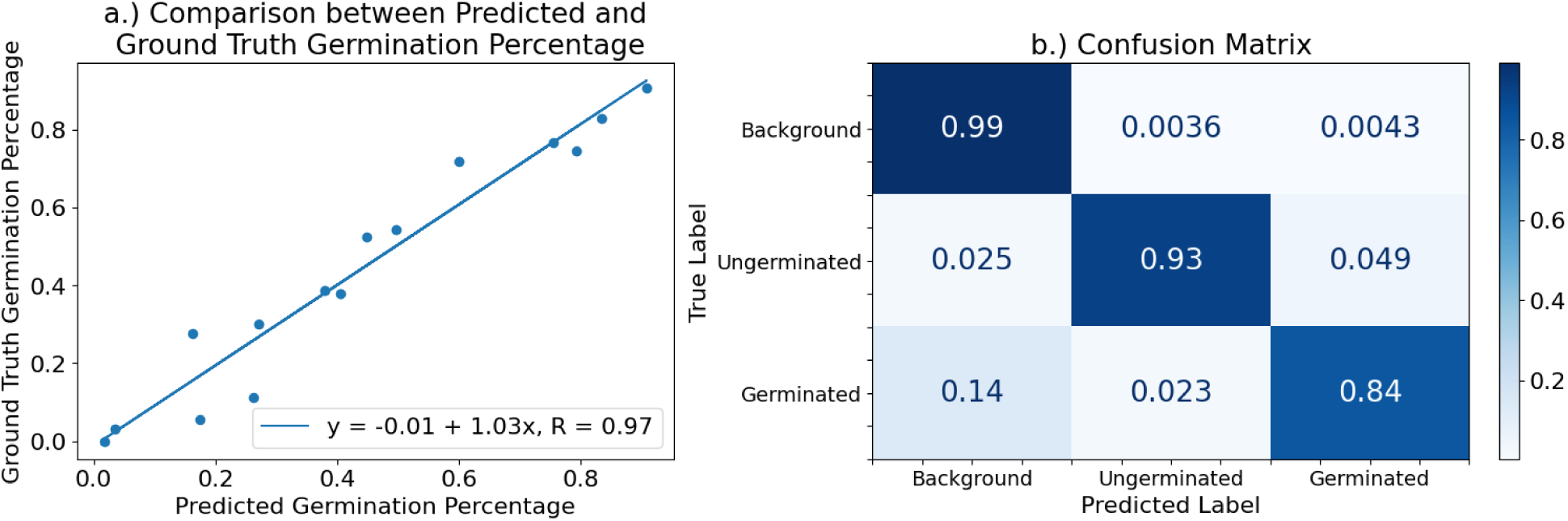
(a) Comparison between ground truth measurements of germination percentage and the predictions for each image in the test set. Percentages were obtained using the test set images annotated by humans for the ground truth and by the machine-learning model for the predictions. Both were run through the measuring software. (b) The confusion matrix compares the predicted labels by the machine learning model to the human-annotated ground truths for each pixel in the test set images, showing the proportion of pixels predicted correctly and incorrectly for each class.

The confusion matrix shows that pixels with the true label of background were almost always correctly labelled (0.99; Figure 3b). The highest proportion of errors (0.14) occurred when germinated pollen was wrongly labelled as background. The accuracy score was 0.9873. The F1-score is above 0.8 for all classes (Table 2), indicating that the model is performing well in terms of both precision and recall. For the germinated pollen, the model has a high precision, but a lower recall. The high precision indicates that the pixels labelled as germinated are likely to be truly germinated, but the lower recall suggests that the model may also predict some of the germinated pixels as ungerminated or background.

**Table 2.** Performance metrics by class for CNN predictions on images in the test set. Precision is the proportion of positive predictions that were truly positive, while recall measures the proportion of the actual positives that were correctly labelled as positive. F1-Score = (2 x Precision x Recall) / (Precision + Recall).

| Metric | Background | Ungerminated | Germinated |
| --- | --- | --- | --- |
| Precision | 0.9928 | 0.8421 | 0.9256 |
| Recall | 0.9959 | 0.8269 | 0.7403 |
| F1-Score | 0.9940 | 0.8269 | 0.8227 |

### 3.2 GP and PL temperature response curves and *T_opt_GP_* and *T_opt_PL_* across the elevational gradient

Both length-based and ratio-based methods were evaluated to quantify pollen germination. Because the length-based approach produced more consistent biological interpretations, only these results are presented here. The ratio-based analyses are provided in the Supplementary Materials. The temperature dependence of GP and PL followed a bell-shaped response across the elevations of pollen collection (Figure 4). The gamma function effectively described the temperature response curves of GP and PL of Douglas-fir (Figure 4). GP showed that populations from high-elevation sites tended to exhibit a lower *T_opt_GP_*, while populations from low-elevation sites exhibited a higher *T_opt_GP_* (Figure 4a). However, *T_opt_PL_* did not show similar trends (Figure 4b). The fitted gamma curves of GP and PL in Douglas-fir at different elevations are available in Supplementary Data Figures S2 and S3.

**Figure 4.**
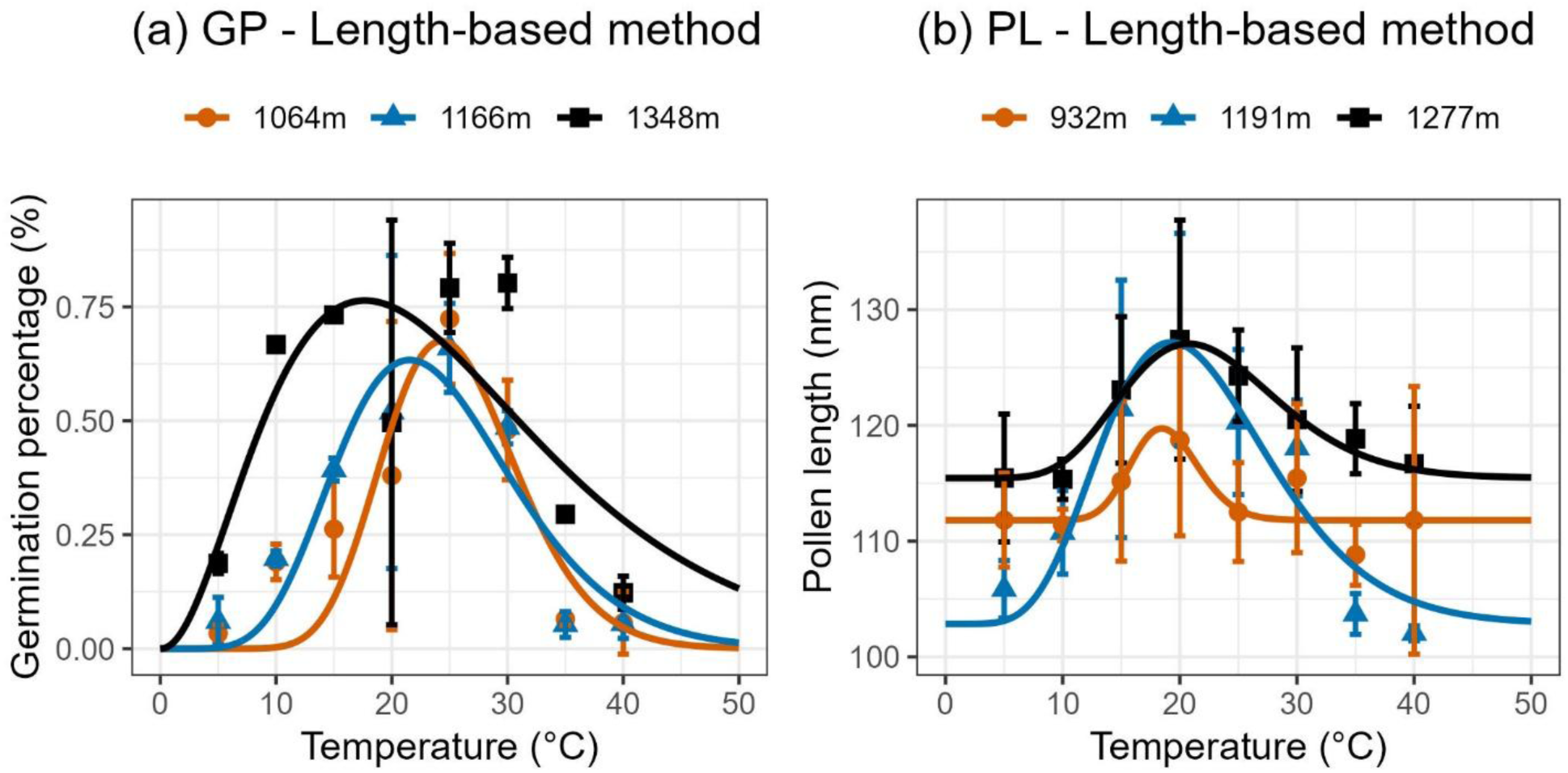
Gamma temperature response curves of pollen germination percentage (a; GP) and pollen length (b; PL) in Douglas-fir from three elevations (low, medium, and high) within the elevational range of collected Douglas-fir pollen.

The mean *T_opt_GP_* and *T_opt_PL_* of collected Douglas-fir pollen were 19.8 and 22.9°C, and were significantly different from PICO, PIEN, and PIPO (*p*<0.001; Table 3). No significant ascending or descending trends were found along the elevational gradient in *T_opt_GP_* and *T_opt_PL_* (*p*=0.136 and 0.522, Figure 5).

**Figure 5.**
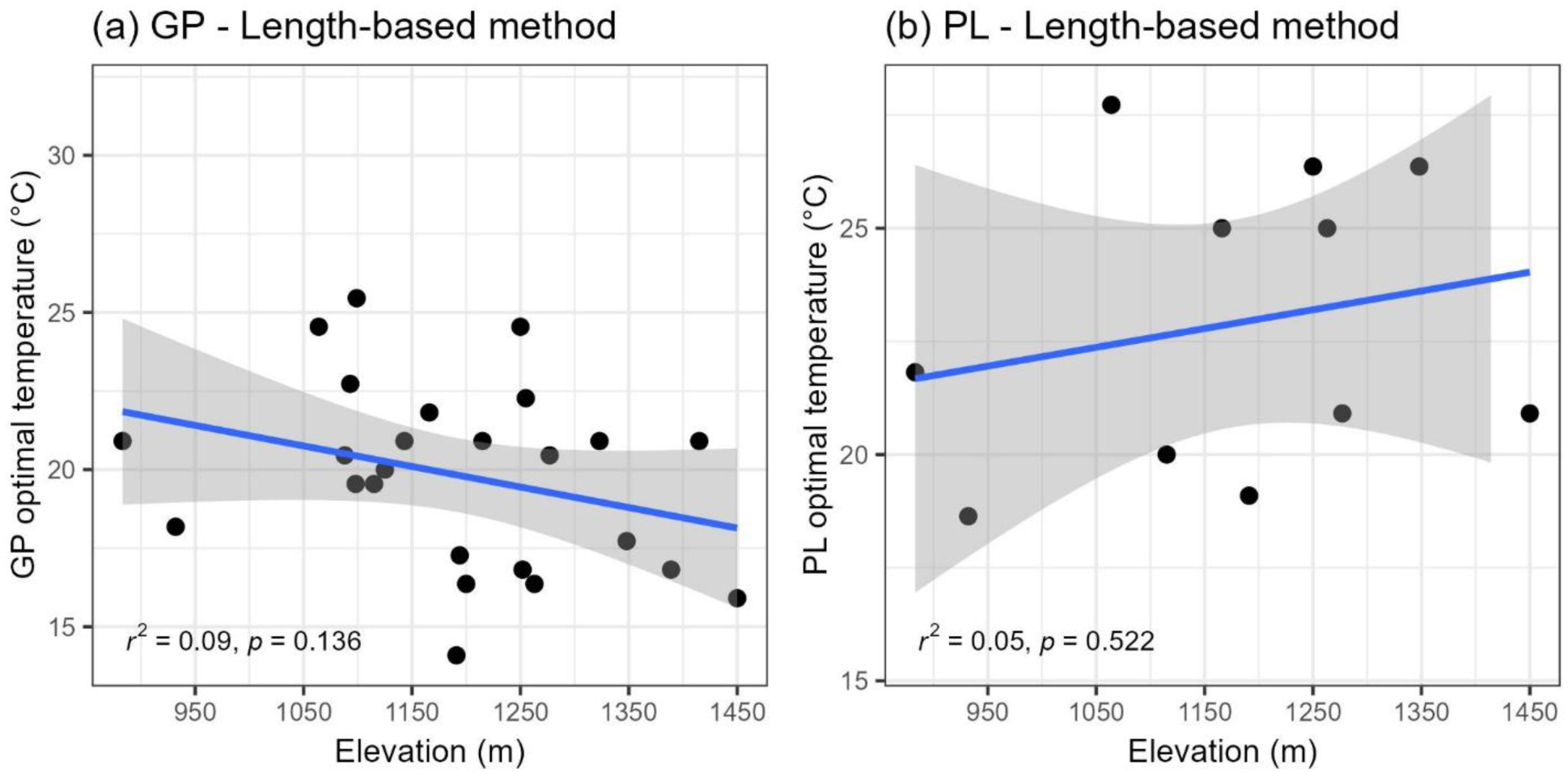
Regression lines of optimal temperatures of pollen germination percentage (a; GP) and pollen length (b; PL) in Douglas-fir across the elevational gradients. The shaded areas are 95 % confidence intervals.

**Table 3.**
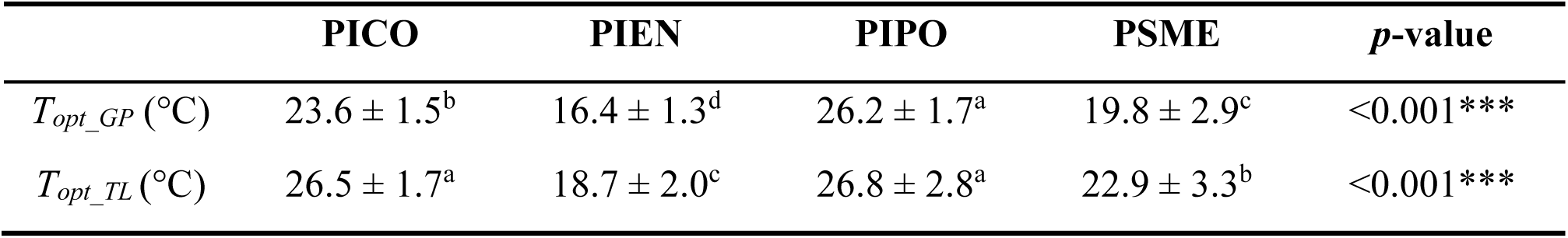
Pollen germination percentage (GP) and pollen length (PL) optimal temperature of *Pseudotsuga menziesii* (PSME) and germination percentage (GP) and pollen tube length (TL) optimal temperature of *Pinus contorta* (PICO), *Picea engelmannii* (PIEN), and *Pinus ponderosa* (PIPO). Different letters indicate significant differences between species (column-wise comparisons) using Tukey’s honest significance test (HSD). \*\*\**p* < 0.001. Data of PICO, PIEN, and PIPO are from Hsu & Kim (2025).

### 3.3 GP and PL vs spring temperature

The *T_opt_GP_* and *T_opt_PL_* of Douglas-fir and the *T_opt_GP_* and *T_opt_TL_* of PICO, PIEN, and PIPO versus the spring mean temperature along the elevational gradient are shown in Figure 6. It showed that each species occupies a unique thermal niche related to its natural habitats.

**Figure 6.**
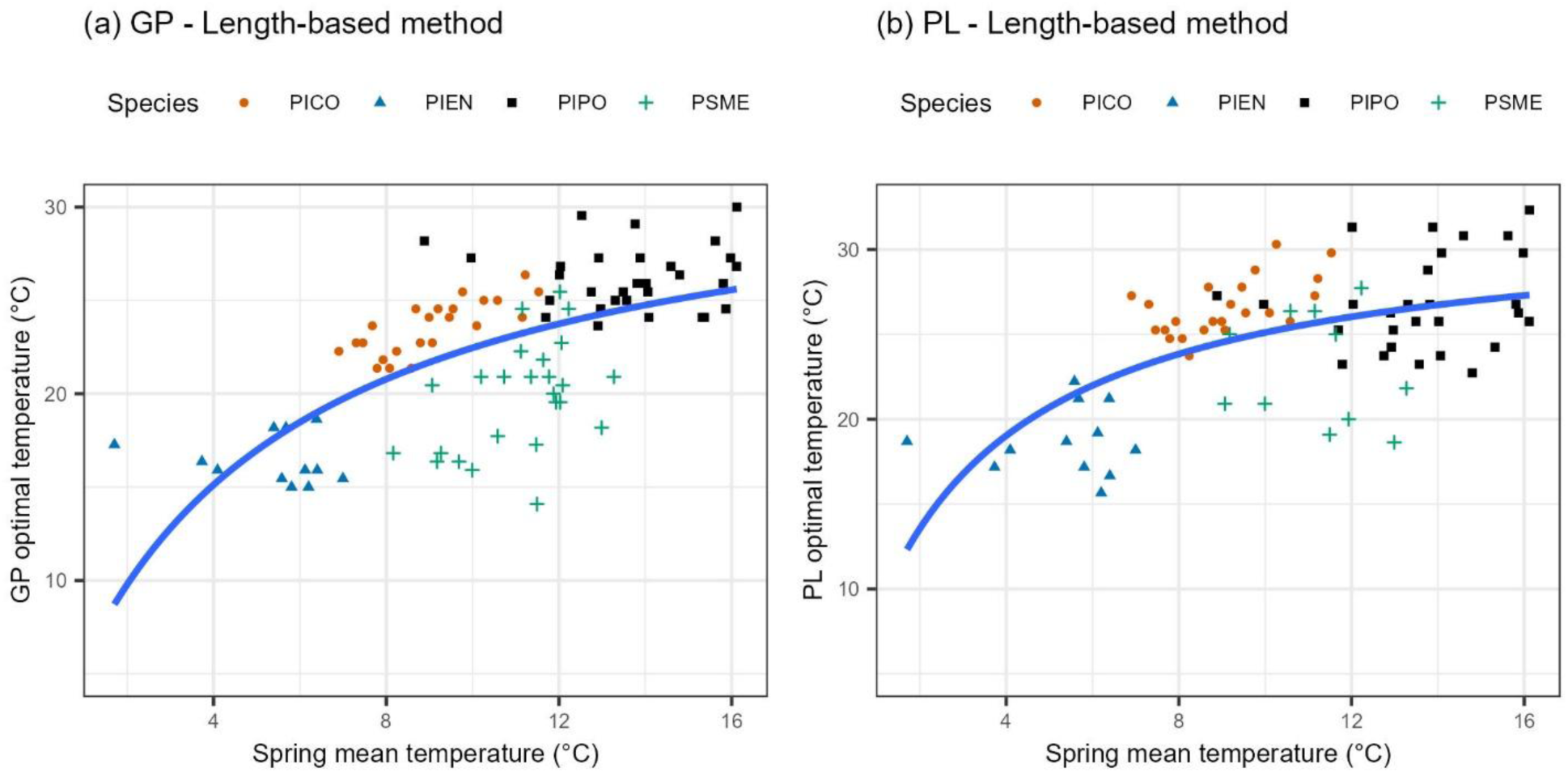
Pollen germination percentage (a; GP) and pollen length (b; PL) optimal temperature of *Pseudotsuga menziesii* (PSME) and GP and TL optimal temperature of *Pinus contorta* (PICO), *Picea engelmannii* (PIEN), and *Pinus ponderosa* (PIPO) versus the spring mean temperature across the elevational gradients. The blue lines are Michaelis–Menten equations. Data of PICO, PIEN, and PIPO are from Hsu & Kim (2025).

### 3.4 GP vs PL optimal temperatures

Populations with higher *T_opt_GP_* would also have higher *T_opt_PL_*; i.e. *T_opt_GP_* and *T_opt_PL_* had a positive relationship (Figure 7). The *T_opt_PL_* was 0.45–8.64 °C higher than the *T_opt_GP_* in PSME. We further tested whether the slope of PSME was significantly different from 1, i.e., whether the increments of *T_opt_GP_* and *T_opt_PL_* were equal. The results showed that they were not (*p* = 0.194).

**Figure 7.**
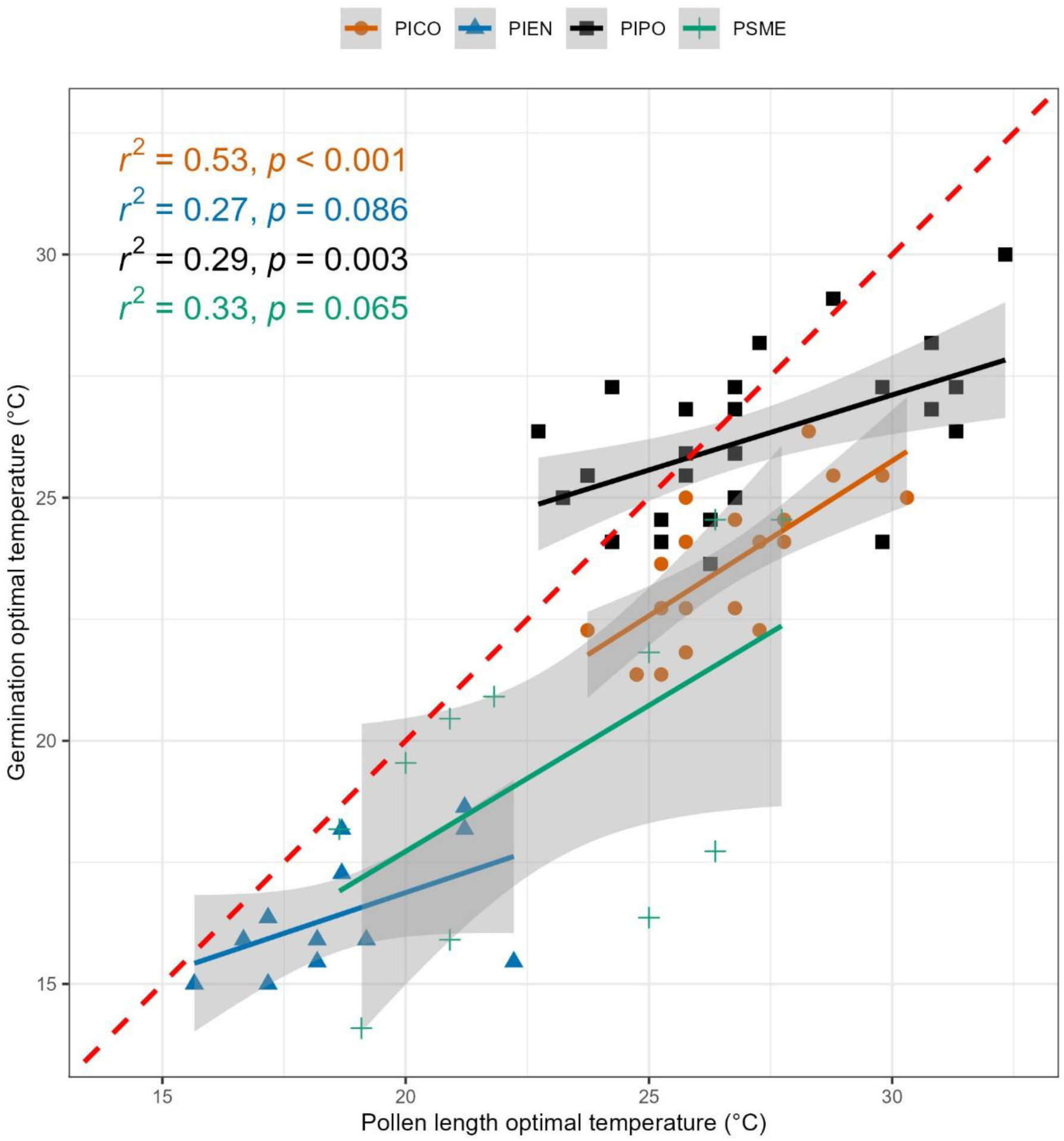
Simple linear regressions between the optimal temperatures of pollen germination percentage (GP) and pollen length (PL) in *Pseudotsuga menziesii* (PSME), and pollen germination percentage and tube length in *Pinus contorta* (PICO), *Picea engelmannii* (PIEN), and *Pinus ponderosa* (PIPO) across the elevational gradients. The red dashed line is the 1:1 temperature ratio. The shaded areas are 95 % confidence intervals. Data of PICO, PIEN, and PIPO are from Hsu & Kim (2025).

## 4. DISCUSSIONS

### 4.1 Performance of deep learning approaches for pollen germination analysis

The CNN developed in this study achieved high segmentation accuracy, with validation IOU improving from 0.699 during initial training to 0.846 after iterative reductions in the learning rate (from 0.5 to 0.005). This improvement highlights the critical importance of optimization strategies in deep-learning applications for biological image analysis (Battiato et al., 2021; Punyasena et al., 2012). Instead of manually decaying the learning rate, using a scheduler or an optimizer that adjusts the learning rate with appropriate hyperparameters could achieve better results. When experimenting with using the Adam optimizer instead of SGD, the model either seemed to be converging too slowly or converging to a model with low accuracy. With more tuning of the learning rate or a different approach to learning rate decay, better results might be achieved. A high initial learning rate enables rapid exploration of the parameter space, whereas gradual reductions allow the model to refine parameters and converge toward improved local optima (Punyasena et al., 2012). These advancements were supported by the PyTorch framework, which utilizes dynamic eager execution to allow for flexible model building and efficient debugging during the iterative optimization process (Arias et al., 2017; Paszke et al., 2019). Additionally, the success of this approach is consistent with other plant phenotyping systems, such as Automated Recognition and Layered Optimization (ARLO), which search for the best combinations of feature weights and resolutions to maximize accuracy (Punyasena et al., 2012). Image analysis with deep learning has been used to develop a variety of high-throughput plant phenotyping methods and is an active area of research (Abebe et al., 2023; Arya et al., 2022). Image segmentation methods have successfully been applied to diverse phenotyping tasks, including root segmentation, leaf counting, wheat spike detection, and maize kernel categorization, as well as many others (Misra et al., 2020; Ubbens & Stavness, 2017; Warman et al., 2021; Yasrab et al., 2019). Data obtained from these high-throughput phenotyping methods can enable the development of tools and technologies in agriculture and plant research, such as process-based plant models, automated pollen detection networks, and precision farming robots (Milioto et al., 2018; Oteros et al., 2020; Ting et al., 2026).

Despite the overall accuracy of the model, classification performance differed substantially among classes. The model distinguished pollen grains from the background with high reliability (IOU: 98.74%) but showed reduced accuracy when separating germinated (IOU: 64.33%) from ungerminated pollen (IOU: 63.14%). Such discrepancies are common in biological imaging tasks where morphological differences between classes are subtle, such as the traditionally difficult task of discriminating between morphologically similar congeneric species like black and white spruce (Punyasena et al., 2012). Background segmentation generally relies on strong contrast differences, whereas identifying germination requires the detection of small protrusions or elongation structures that may vary in orientation and visibility (Gonçalves et al., 2016; Toda et al., 2021). For species like Douglas-fir, this is further complicated by its unique two-step germination process, where grains first undergo significant elongation before producing a distinct pollen tube (Dumont-BéBoux & Von Aderkas, 1997; Fernando et al., 2005). This atypical development means that early elongation may not be immediately distinguishable from the turgid, oval state of ungerminated grains, representing a significant challenge for automated detection (Arias et al., 2017). In our study, because the germinated and ungerminated pollen look dissimilar in shape and color, they were labeled separately when training the model, but both germinated and ungerminated pollen were considered as one class, and distinguishing them solely through measurement might be as accurate and simpler for the annotators. For some of the images in the test set, the per-class accuracies were much lower than in other images. This variance is not immediately clear from the average. Taking a closer look at these images could reveal how accuracy could be improved. For example, if these kinds of images are underrepresented in the training set, more representative images could be added.

The model also tended to underestimate germination percentages relative to manual counts. This bias likely reflects systematic underestimation of early pollen elongation. In many standard *in vitro* germination assays, a pollen grain is considered germinated when the tube length exceeds the diameter of the grain (Bohrerova et al., 2009; Cram & Lindquist, 1984; Hsu & Kim, 2025; Pham et al., 2015; Pio et al., 2004). Small measurement errors near this threshold can lead to misclassification (Hsu & Kim, 2025). In Douglas-fir specifically, standard protocols often categorize germination into classes based on whether an elongating grain has reached twice its original diameter (Webber & Bonnet-Masimbert, 1993; Webber & Painter, 1996). Consistent with this interpretation, the most frequent error observed in the test dataset occurred when germinated pollen grains were incorrectly labeled as background. These errors suggest that the model had relatively high precision (0.925) but lower recall (0.740), a phenomenon also observed in StomataCounter and other deep-learning models where existing objects are “missed” due to blur or artifacts, leading to undercounting (Fetter et al., 2019). The high precision indicates that the model is highly reliable when it identifies a grain as germinated; however, the lower recall reveals that it “misses” a significant portion of truly germinated grains.

Several factors may contribute to these false-negative errors. Pollen grains often occur in clusters or clumps, which can complicate segmentation and have been noted to potentially affect germination rates *in vitro* (Duffield, 1954). Microscopic images may also contain background artifacts, such as debris or air bubbles, that reduce contrast between grains and the surrounding medium (Gonçalves et al., 2016; Khanzhina et al., 2018). Recent large-scale datasets like Pollen13K specifically include classes for “debris” to help models distinguish between artifacts and actual pollen (Battiato et al., 2021). Therefore, increasing the diversity of the training dataset, especially by including more examples of clumped pollen grains or low-contrast images, may improve model generalization capability (Battiato et al., 2021; Khanzhina et al., 2018).

Future improvements may involve simplifying the classification task. Instead of separating germinated and ungerminated grains during segmentation, it may be advantageous to detect all pollen grains as a single class and determine germination status using subsequent morphological measurements (Fetter et al., 2019; Toda et al., 2021). Integrating attention gates (AGs), such as those in the Attention U-Net, could further improve results by suppressing irrelevant background regions and highlighting salient features like early elongation protrusions (Arias et al., 2017; Oktay et al., 2018). Improvements in image acquisition, such as the use of Environmental Scanning Electron Microscopy (ESEM) or confocal microscopy to resolve fine surface detail and three-dimensional structures, could also enhance model performance (Colangeli & Owens, 1990; Dawkins & Owens, 1993; Losada & Leslie, 2018; Nel et al., 2005). For example, focal stacking or three-dimensional z-stack imaging provides clearer visualization of pollen tubes by combining information from multiple focal planes, improving the detection of small, out-of-focus elongation structures (Gonçalves et al., 2016; Khanzhina et al., 2018; Punyasena et al., 2012).

Automated phenotyping tools, such as the model presented in our study, have increasing importance in plant reproductive biology because they enable high-throughput quantification of traits that are otherwise labor-intensive to measure (Fetter et al., 2019; Gonçalves et al., 2016; Hsu & Kim, 2025; Toda et al., 2021; Tushabe & Rosbakh, 2021). By accelerating pollen germination and growth analysis, these tools facilitate extensive comparative studies on reproductive traits, the “weak link” in plant climate responses, across varying elevations and climatic conditions (Barnabás et al., 2008; Flores-Rentería et al., 2018; Gonçalves et al., 2016; Hebbar et al., 2018; Hedhly et al., 2009; Hsu & Kim, 2025; Kakani et al., 2002; Rosbakh & Poschlod, 2016).

### 4.2 Temperature sensitivity of pollen germination and growth

Temperature is a primary environmental factor controlling pollen germination and pollen tube growth, influencing both the speed of the process and the ability of the male gamete to reach the ovule (Hebbar et al., 2018; Hsu & Kim, 2025; Kakani et al., 2002; Steinacher & Wagner, 2012). Both processes typically exhibit bell-shaped responses to temperature, characterized by clearly defined minimum, optimum, and maximum thresholds (Hebbar et al., 2018; Rosbakh & Poschlod, 2016; Thakur et al., 2010), and with performance increasing up to an optimal temperature before declining at supraoptimal levels (Hsu & Kim, 2025; Kakani et al., 2002; Steinacher & Wagner, 2012). In our study, temperature response curves for GP and PL were well described using gamma functions, indicating that Douglas-fir pollen exhibits a clear thermal optimum for reproductive performance. The use of the gamma function is consistent with conifer studies, as it provides additional flexibility to model the specific ascending and decay rates around the optimum (Hsu & Kim, 2025).

The estimated optimal temperatures for Douglas-fir fell within a moderate range of approximately 19–23°C. These values are consistent with established *in vitro* observations for the species, which place the optimum between 20–25°C (Ho, 1968). These ranges are broadly consistent with findings that conifer pollen generally achieves maximal germination at intermediate temperatures (Hsu & Kim, 2025; Varis et al., 2011), though Douglas-fir optima appear slightly lower than some other members of the *Pinaceae*. For example, studies on several *Pinus* species found optimal germination closer to 30–32°C(Flores-Rentería et al., 2018; Nel et al., 2005) and 23.6 and 26.2°C for *Pinus contorta* and *Pinus ponderosa* (Hsu & Kim, 2025), while higher than *Picea Engelmannii* peaks at 16.4°C (Hsu & Kim, 2025). Notably, it is a recognized pattern in conifers that the optimal temperature for growth (length) is generally higher than the optimal temperature for germination, likely because tubes must adapt to the higher temperatures that occur later in the season following the initial pollination event (Hsu & Kim, 2025). The relatively narrow and moderate thermal range observed for Douglas-fir suggests that its reproductive performance is highly sensitive to shifts in environmental temperature during the critical pollination window (Flores-Rentería et al., 2018; Hedhly et al., 2009; Hsu & Kim, 2025).

Temperature sensitivity during the progamic phase, the period between pollination and fertilization, is critical because this sequence of processes must be highly coordinated for fertilization to succeed (Hedhly et al., 2009; Karabiyik, 2022; Steinacher & Wagner, 2012). In Douglas-fir, this phase is unique and extended, involving an initial period where the grain hydrates and elongates within the micropylar canal, followed by a second phase of tube formation approximately one week before fertilization (Dumont-BéBoux & Von Aderkas, 1997; Fernando et al., 2005; Owens et al., 1998). Environmental conditions during this stage can influence reproductive success even when pollen production and dispersal remain unaffected (Flores-Rentería et al., 2018; Hebbar et al., 2018; Hedhly et al., 2009; Hsu & Kim, 2025). This potential for developmental asynchrony means that even a slight increase in mean spring temperatures (1.4–3°C) can lead to erratic fruit set and reproductive failure (Hedhly et al., 2009). Because *in vitro* pollen germination tests are strong predictors of total seed and filled seed percentages (Moody & Jett, 1990; Webber & Bonnet-Masimbert, 1993), our results illustrate that temperature-induced failures during this phase can be a major limitation on tree reproduction under a rapidly shifting climate (Hsu & Kim, 2025; Webber, 1987).

### 4.3 Elevational variation and local adaptation

No significant relationship was detected between elevation and the optimal temperature for pollen germination in Douglas-fir populations (*p* = 0.136). Although visual inspection of the fitted response curves suggested that some high-elevation populations exhibited lower optimal temperature than certain low-elevation populations, this pattern was inconsistent across the elevational gradient and should be interpreted with caution. Consequently, our data do not provide strong evidence for an elevational cline in pollen germination temperature optima within Douglas-fir.

The lack of a significant elevational trend contrasts with expectations based on local adaptation theory, which predicts that populations occupying cooler environments may evolve lower thermal optima for reproductive processes (Hedhly et al., 2009; Hsu & Kim, 2025; Rosbakh & Poschlod, 2016). Previous studies have shown that pollen germination and tube growth temperatures can vary among populations and species along climatic gradients, reflecting adaptation to local thermal environments (Hsu & Kim, 2025; Rosbakh & Poschlod, 2016). However, our results suggest that any such differentiation in Douglas-fir is either weak, obscured by environmental heterogeneity, or occurs at a scale not captured by elevation alone.

The absence of statistical significance across the entire gradient for Douglas-fir may reflect the influence of microclimatic variability. Elevation serves only as a proxy for temperature, and the actual thermal conditions experienced by pollen may vary substantially depending on local factors such as slope aspect, canopy structure, and exposure to net radiation (Körner, 2007; Pigott & Huntley, 1981; Rosbakh & Poschlod, 2016). These microenvironmental differences can decouple the environment that individuals actually experience from the predictors used in broad analysis. Such microclimatic decoupling, along with potentially narrow geographical sampling ranges or confounding factors like regional moisture regimes, may mask broader elevational patterns in pollen performance (Hsu & Kim, 2025; Körner, 2007). Furthermore, developmental stages for Douglas-fir at higher elevations can be up to a month later than those at lower elevations, potentially exposing these populations to different weather patterns that obscure broad elevational trends (Fernando et al., 1997; Rosbakh & Poschlod, 2016). Such microenvironmental differences, combined with the high degree of phenotypic plasticity often seen in plant reproductive traits, can mask larger-scale patterns of local climatic adaptation (Hedhly et al., 2009).

### 4.4 Distinct thermal optima for germination and pollen growth

A key result of this study was the difference between optimal temperatures for pollen germination and pollen elongation. The optimal temperature for pollen elongation was consistently higher than the optimal temperature for germination by 0.45-8.64°C. This pattern has been observed in multiple plant taxa and likely reflects differences in the physiological processes that tube growth requires higher thermal energy than initial germination (Hebbar et al., 2018; Hsu & Kim, 2025; Parantainen & Pasonen, 2004). Pollen germination represents the activation of the pollen grain and involves hydration, metabolic reactivation, and initiation of polarized growth (Dafni & Firmage, 2000; Hedhly et al., 2009; Taylor & Hepler, 1997; Webber & Painter, 1996). In contrast, pollen tube elongation is a highly dynamic cellular growth process that requires rapid synthesis of cell wall components and continuous cytoplasmic transport (Çetinbaş-Genç et al., 2019; Dawkins & Owens, 1993; Fernando et al., 2005; Taylor & Hepler, 1997). Because these processes involve intense metabolic activity, tube growth often accelerates at temperatures slightly higher than those optimal for germination (Hedhly et al., 2004, 2009; Hsu & Kim, 2025; Steinacher & Wagner, 2012; Thakur et al., 2010). Furthermore, the transition from autotrophic growth (relying on internal reserves) to heterotrophic growth (utilizing external nutrients) may further explain these diverging thermal sensitivities (Hedhly et al., 2004; Tushabe & Rosbakh, 2021).

The positive relationship between *T_opt_GP_* and *T_opt_PL_* suggests that populations adapted to warmer germination environments also support faster pollen growth at higher temperatures (Hsu & Kim, 2025). However, the increments between these optima were not equivalent, indicating that germination and growth respond differently to temperature (Hsu & Kim, 2025). This result supports the hypothesis that these stages of the progamic phase are at least partially regulated by independent physiological mechanisms and genetic control and subjected to different environmental filters (Hsu & Kim, 2025; Parantainen & Pasonen, 2004).

### 4.5 Biological specificity of Douglas-fir pollen development

Interpretation of these results must account for the unique two-step germination process of Douglas-fir pollen, which distinguishes it from many other conifer species (Dumont-BéBoux & Von Aderkas, 1997; Fernando et al., 2005; Ho, 1968; Webber, 1987). Unlike many conifer species that form a distinct pollen tube immediately after germination, Douglas-fir pollen initially undergoes elongation of the entire grain before a clearly defined tube develops. This elongation results in a cigar-shaped pollen grain during early development. Because of this specific morphology, changes in pollen length and changes in the length-to-diameter ratio do not necessarily occur simultaneously or linearly as they might in species with traditional tubes (Ho, 1968). The elongation of the grain itself fundamentally alters its geometry, reaching lengths of approximately 200–500 µm before a distinct, narrow pollen tube finally develops (Fernando et al., 1997, 2005). The length-based approach better captured the characteristic elongation of Douglas-fir pollen because they directly quantify the physical expansion of the grain, which is the primary morphological change occurring during the first week of germination (Dumont-BéBoux & Von Aderkas, 1997; Ho, 1968) and therefore provided the most biologically meaningful estimates of thermal performance. This unique developmental pattern suggests that germination in Douglas-fir is a more prolonged and complex process than in many other *Pinaceae* (Dumont- BéBoux & Von Aderkas, 1997; Fernando et al., 2005).

### 4.6 Implications for forest reproduction under climate change

Understanding the temperature sensitivity of pollen germination and growth is essential for predicting how plant reproductive processes may respond to climate change (Hebbar et al., 2018; Hedhly et al., 2009; Hsu & Kim, 2025). The optimal temperatures identified in this study (approximately 19.8–22.9°C) suggest that Douglas-fir pollen performance occurs within a relatively narrow thermal window, a finding consistent with the intermediate temperature requirements typically observed in conifer species (Hsu & Kim, 2025). Projected increases in spring temperatures across many regions of western North America, estimated to rise between 1.4°C and 4.4°C by the end of the century, may shift environmental conditions beyond these optimal ranges (Flores-Rentería et al., 2018; Hsu & Kim, 2025). Although moderate warming could potentially accelerate pollen elongation or tube growth, excessive temperatures may lead to irreversible functional disruptions, such as abnormal shapes or polarity issues (Ho, 1968; Steinacher & Wagner, 2012), reduce germination percentages (Barnabás et al., 2008; Hsu & Kim, 2025), or disrupt the synchronization between male and female reproductive structures and lead to complete pollen sterility (Flores-Rentería et al., 2018; Hedhly et al., 2009). Reproductive success in conifers depends on precise timing between pollen germination, pollen tube growth, and ovule receptivity (Fernando et al., 2005; Hedhly et al., 2009; Steinacher & Wagner, 2012). If warming temperatures accelerate pollen development but also shorten the period of ovule receptivity, fertilization success may decline (Hedhly et al., 2009). Such developmental asynchrony has been identified as an important factor influencing seed set in many species (Hedhly et al., 2004, 2009). Consequently, shifts in the thermal environment during the pollination period may influence reproductive outcomes even when vegetative growth remains relatively unaffected (Flores-Rentería et al., 2018; Hedhly et al., 2009; Hsu & Kim, 2025; Sorkheh et al., 2018).

Variation in pollen performance across temperature ranges may also influence genetic diversity within populations (Hsu & Kim, 2025). At suboptimal temperatures, reduced variation in pollen growth rates may allow a broader range of genotypes to successfully fertilize ovules, potentially increasing genetic diversity. In contrast, under optimal conditions, faster-growing pollen may dominate fertilization, reducing diversity. These dynamics suggest that temperature not only affects reproductive success but may also shape the genetic structure of future populations. Furthermore, incorporating pollen thermal responses into species distribution models could improve predictions of plant responses to climate change. Integrating such physiological thresholds into models may enhance our ability to predict range shifts and population persistence under future climate scenarios (Hsu & Kim, 2025). Therefore, understanding the physiological limits of pollen performance is critical for predicting future patterns of forest regeneration and for developing management strategies that maintain reproductive success under changing climatic conditions (Hsu & Kim, 2025; Rosbakh & Poschlod, 2016; Webber & Painter, 1996).

## 5. CONCLUSIONS

CNN-based Deep Learning enabled accurate, high-throughput analysis of Douglas-fir pollen, achieving strong segmentation performance but highlighting limitations in distinguishing early germination stages. These results suggest that simplifying classification and measuring pollen size and shape after detection may improve robustness. Pollen germination and tube elongation showed clear, unimodal temperature-dependent responses with varying optimal temperatures, indicating that these processes are partly controlled by different mechanisms. The species’ two-step germination process further highlights the need to match analytical methods with its biology. Although elevational trends were not statistically significant, variation in optimal temperatures is consistent with possible local adaptation. Overall, pollen performance occurs within a narrow temperature range, suggesting high sensitivity to climate warming. Together with previous findings across conifer species, our results indicate that species-specific reproductive thermal niches are closely associated with local climatic conditions. These findings improve our understanding of the thermal sensitivity of conifer reproduction and provide a scalable framework for assessing the impacts of climate warming on forest regeneration.

## SUPPORTING INFORMATION

Figure S1. Training and validation IOU for stochastic gradient descent during the training process. Xavier Normal initialization was used, and the learning rate was set to 0.5 for the first 200 epochs, then decayed to 0.1 for 30 epochs, 0.5 for 25 epochs, and 0.005 for 30 epochs.

Figure S2. Gamma temperature response curves of length-based pollen germination percentage (GP) in *Pseudotsuga menziesii* (PSME) from different elevations.

Figure S3. Gamma temperature response curves of length-based pollen length (PL) in *Pseudotsuga menziesii* (PSME) from different elevations.

Figure S4. Gamma temperature response curves of ratio-based pollen germination percentage (a; GP) and pollen length (b; PL) in Douglas-fir from three elevations (low, medium, and high) within the elevational range of collected Douglas-fir pollen.

Figure S5. Gamma temperature response curves of ratio-based pollen germination percentage (GP) in *Pseudotsuga menziesii* (PSME) from different elevations.

Figure S6. Gamma temperature response curves of ratio-based pollen length (PL) in *Pseudotsuga menziesii* (PSME) from different elevations.

Figure S7. Regression lines of optimal temperatures of ratio-based pollen germination percentage (a; GP) and pollen length (b; PL) in Douglas-fir across the elevational gradients. The shaded areas are 95 % confidence intervals.

Figure S8. Ratio-based pollen germination percentage (a; GP) and pollen length (b; PL) optimal temperature of *Pseudotsuga menziesii* (PSME) and GP and TL optimal temperature of *Pinus contorta* (PICO), *Picea engelmannii* (PIEN), and *Pinus ponderosa* (PIPO) versus the spring mean temperature across the elevational gradients. The blue lines are Michaelis–Menten equations. Data of PICO, PIEN, and PIPO are from Hsu & Kim (2025).

Figure S9. Simple linear regressions between the optimal temperatures of pollen germination percentage (GP) and pollen length (PL) in *Pseudotsuga menziesii* (PSME), and pollen germination percentage and tube length in *Pinus contorta* (PICO), *Picea engelmannii* (PIEN), and *Pinus ponderosa* (PIPO) across the elevational gradients. The red dashed line is the 1:1 temperature ratio. The shaded areas are 95 % confidence intervals. Data of PICO, PIEN, and PIPO are from Hsu & Kim (2025).

Table S1. Ratio-based pollen germination percentage (GP) and pollen length (PL) optimal temperature of *Pseudotsuga menziesii* (PSME) and germination percentage (GP) and pollen tube length (TL) optimal temperature of *Pinus contorta* (PICO), *Picea engelmannii* (PIEN), and *Pinus ponderosa* (PIPO). Different letters indicate significant differences between species (column- wise comparisons) by using Tukey’s honest significance test (HSD). ***p < 0.001. Data of PICO, PIEN, and PIPO are from Hsu & Kim (2025).

## AUTHOR CONTRIBUTIONS

Hsin-Wu Hsu: Writing – original draft, Writing – review & editing, Visualization, Software, Methodology, Investigation, Formal analysis, Conceptualization.

Sabrina Zerrade: Writing – original draft, Writing – review & editing, Visualization, Software, Methodology, Formal analysis.

Soo-Hyung Kim: Writing – review & editing, Supervision, Methodology, Funding acquisition, Conceptualization.

## DATA AVAILABILITY

Data is available on Dryad. DOI: 10.5061/dryad.djh9w0wgg

## ACKNOWLEDGEMENTS

We thank undergraduate students Reve Hansen, Abigail Lovell, Sriram Parasurama, Sarah Shafer, Shannon Thoele and Leah Valentine at the School of Environmental and Forest Sciences, University of Washington, and volunteers Susan Bicais, Caitlin Maraist, Colton Stokes and Stephanie Woods for helping with conducting pollen viability tests, taking pollen images, counting pollen germination or measuring pollen tube length during the COVID-19 pandemic. We thank undergraduate students who took ESRM499 courses from winter 2022 to spring 2023 at the School of Environmental and Forest Sciences, University of Washington, for counting pollen germination and measuring pollen tube length.

## APPENDIX

**Table A.**
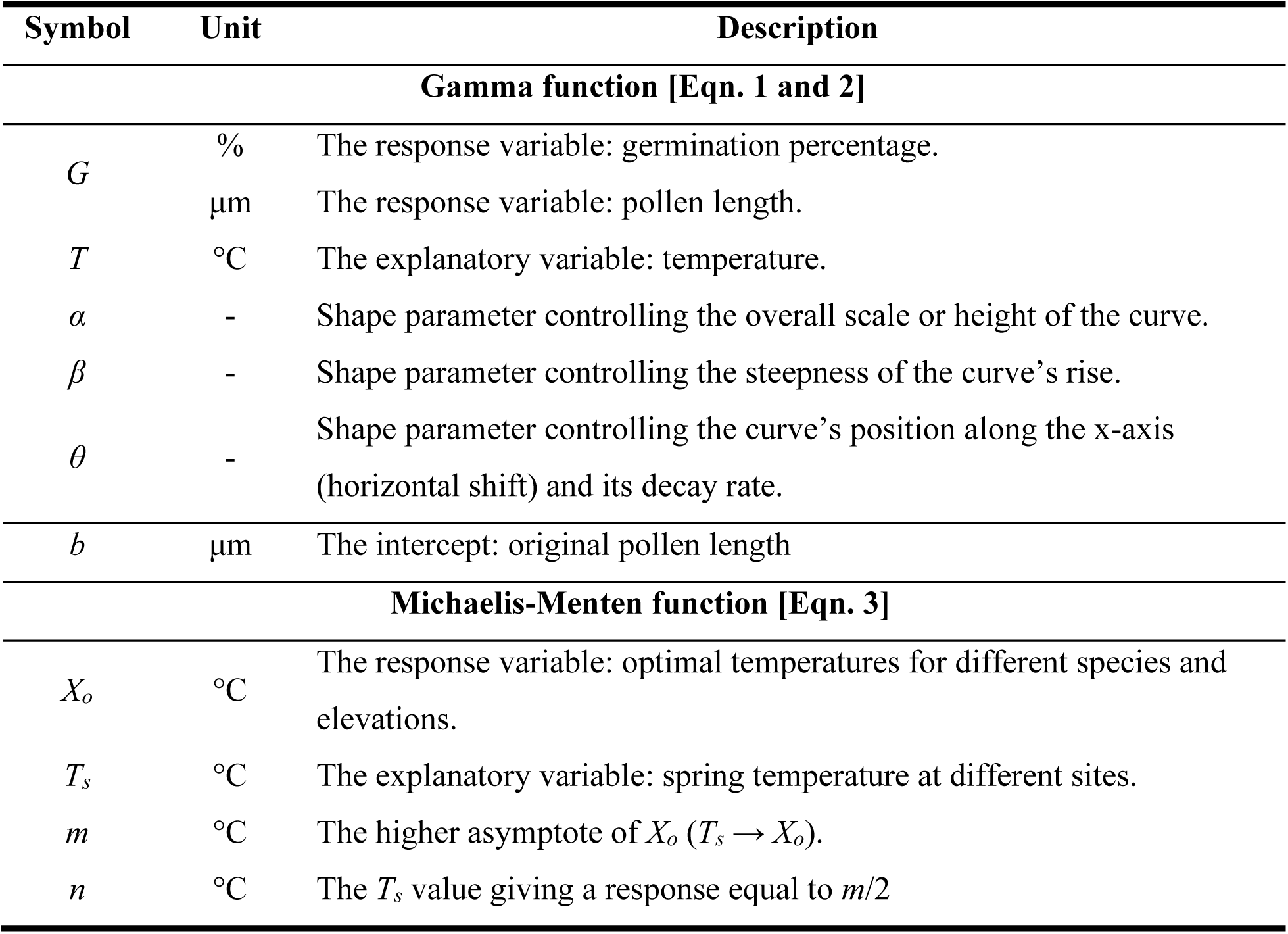
Variables and parameters used in the Gaussian, Gamma, and Michaelis-Menten functions and their descriptions.

