## Supplementary materials for "Temperature dependence of pollen germination in Douglas-fir (*Pseudotsuga menziesii*): A machine learning-based detection of pollen viability from microscopic images"

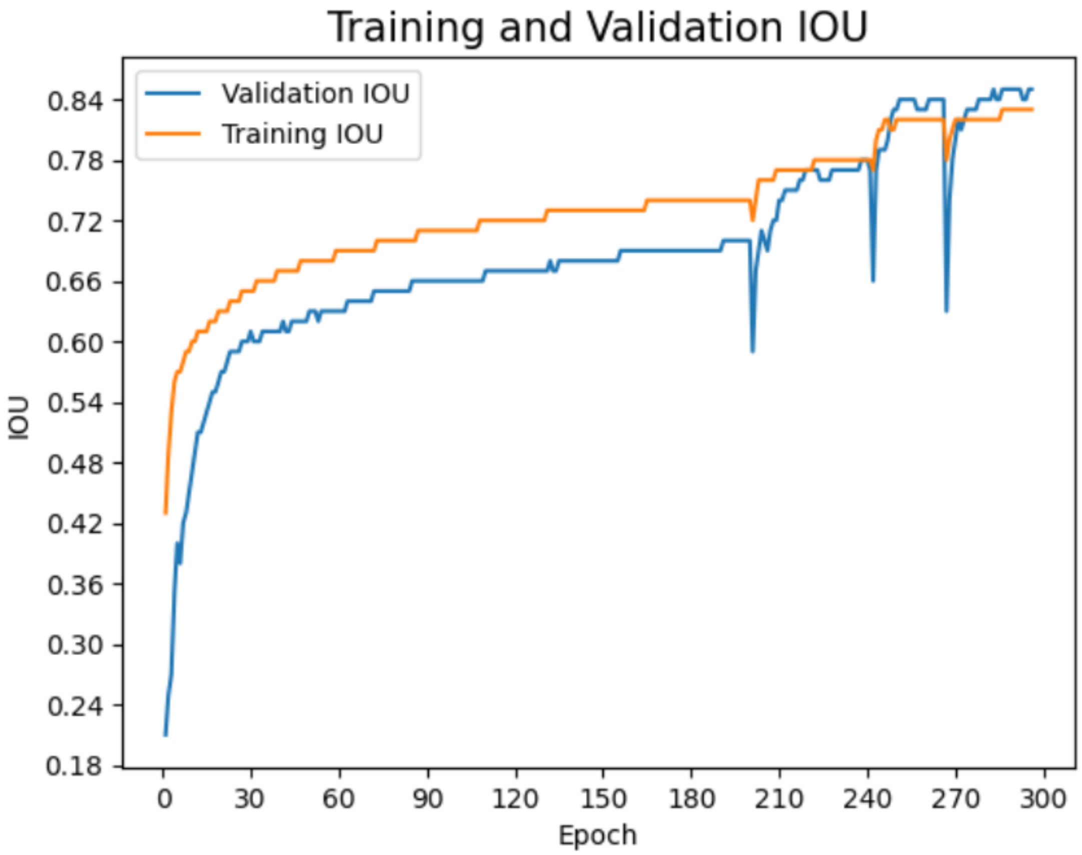


Figure S1. Training and validation IOU for stochastic gradient descent during the training process. Xavier Normal initialization was used, and the learning rate was set to 0.5 for the first 200 epochs, then decayed to 0.1 for 30 epochs, 0.5 for 25 epochs, and 0.005 for 30 epochs.

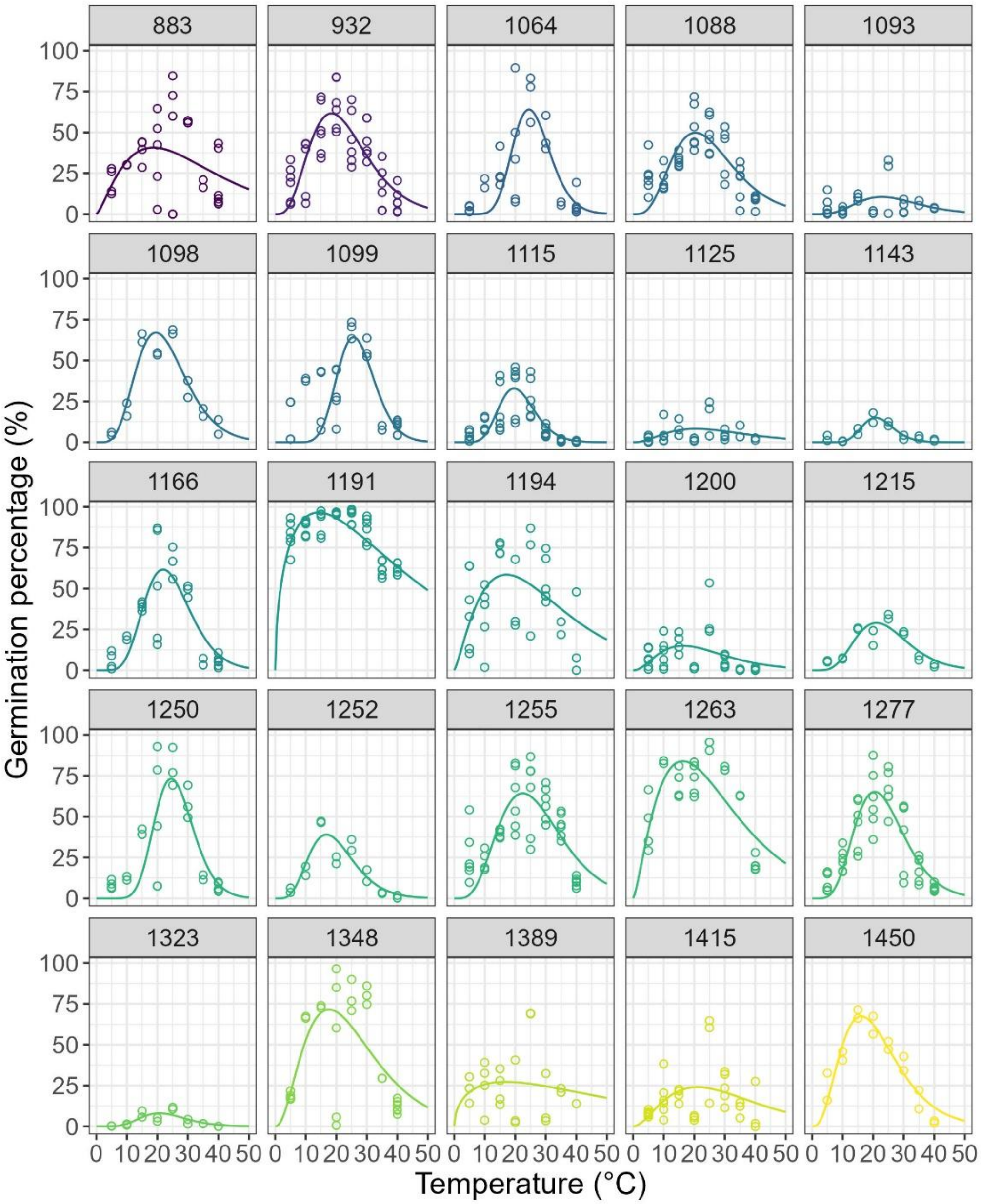


Figure S2. Gamma temperature response curves of length-based pollen germination percentage (GP) in *Pseudotsuga menziesii* (PSME) from different elevations.

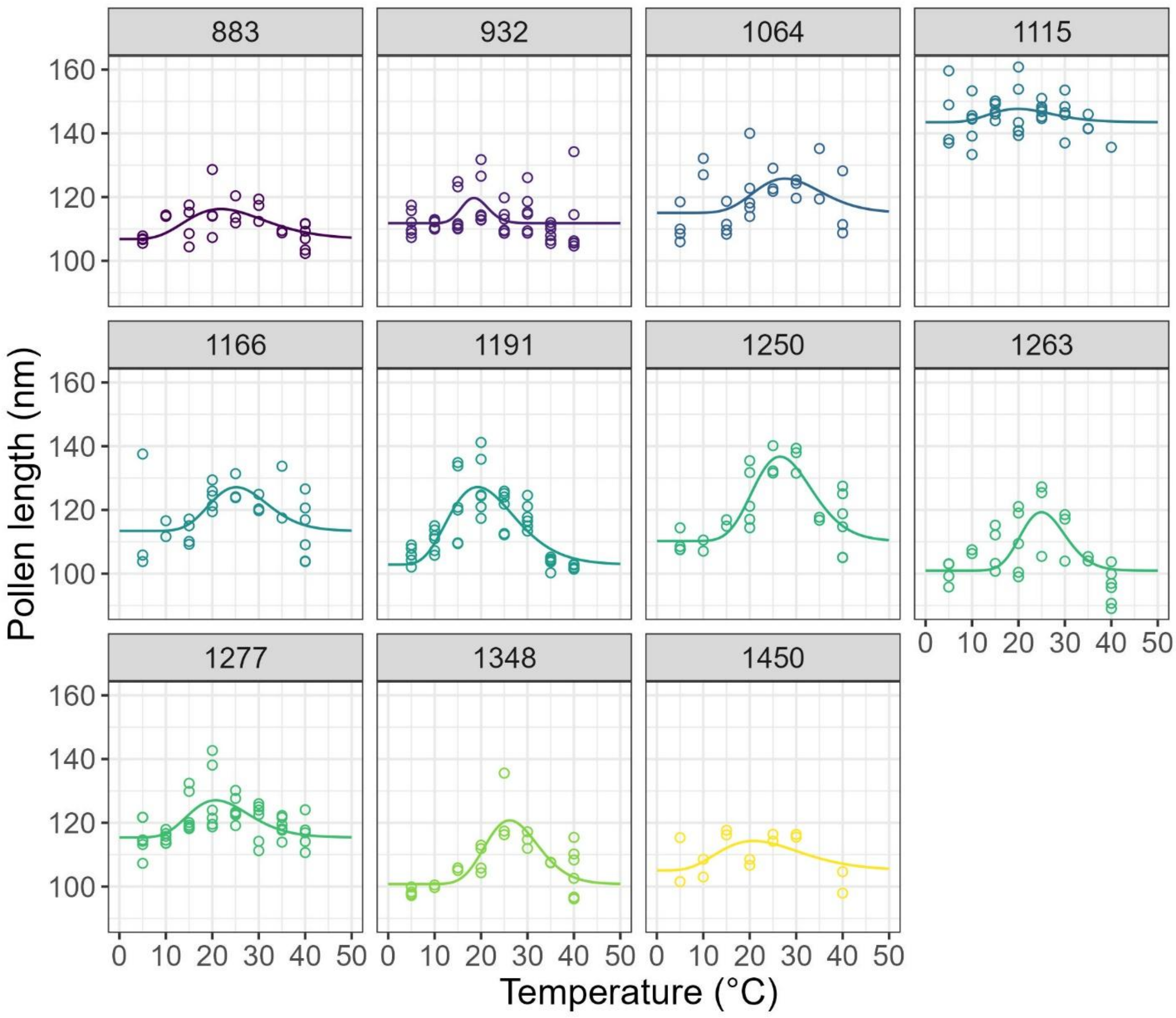


Figure S3. Gamma temperature response curves of length-based pollen length (PL) in *Pseudotsuga menziesii* (PSME) from different elevations.

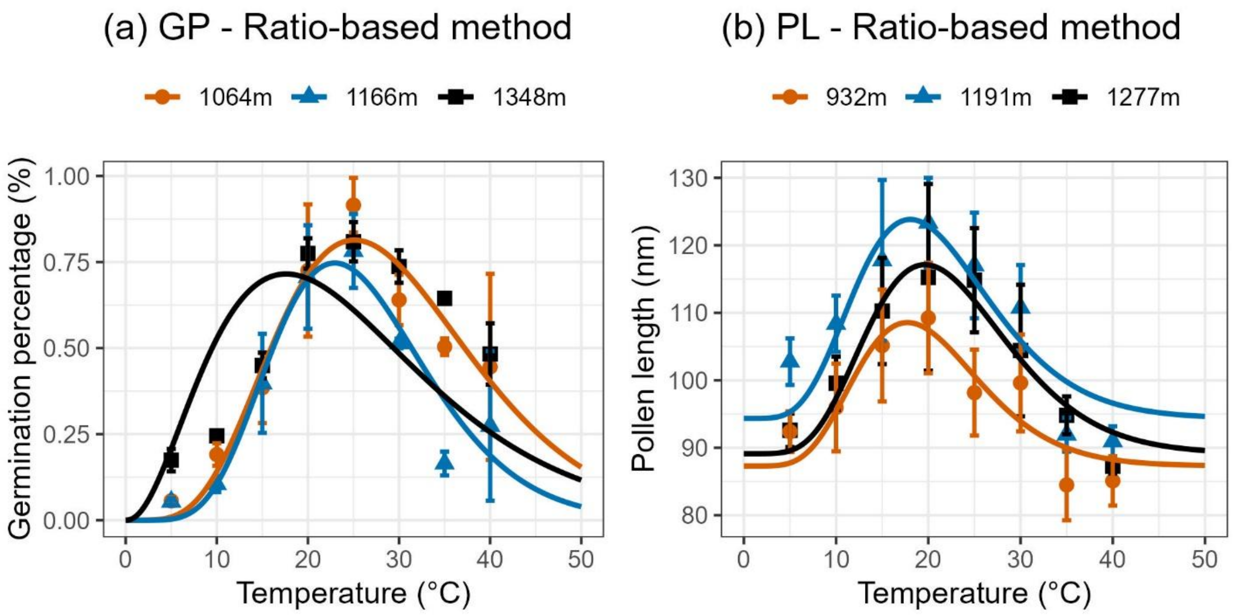


Figure S4. Gamma temperature response curves of ratio-based pollen germination percentage (a; GP) and pollen length (b; PL) in Douglas-fir from three elevations (low, medium, and high) within the elevational range of collected Douglas-fir pollen.

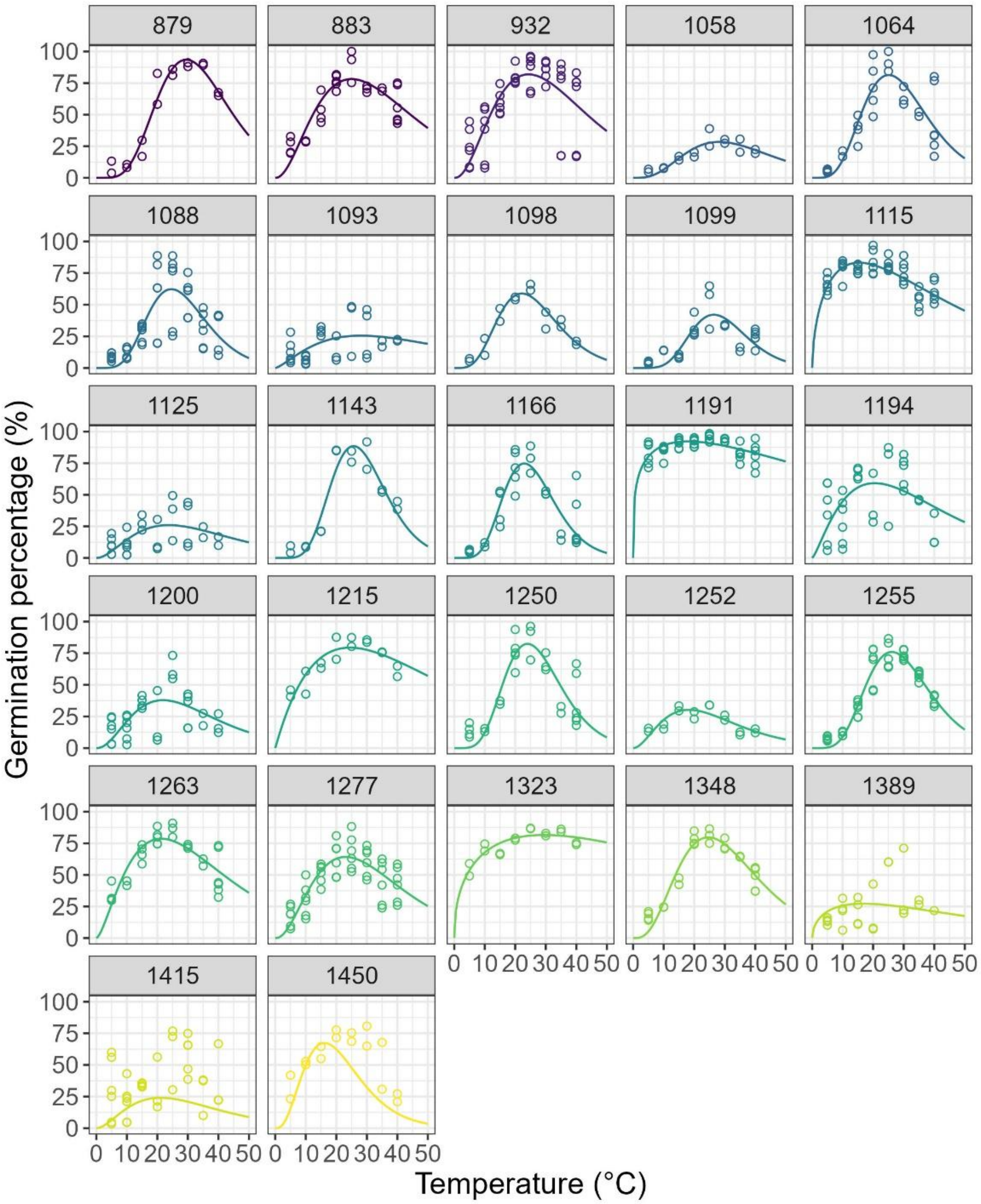


Figure S5. Gamma temperature response curves of ratio-based pollen germination percentage (GP) in *Pseudotsuga menziesii* (PSME) from different elevations.

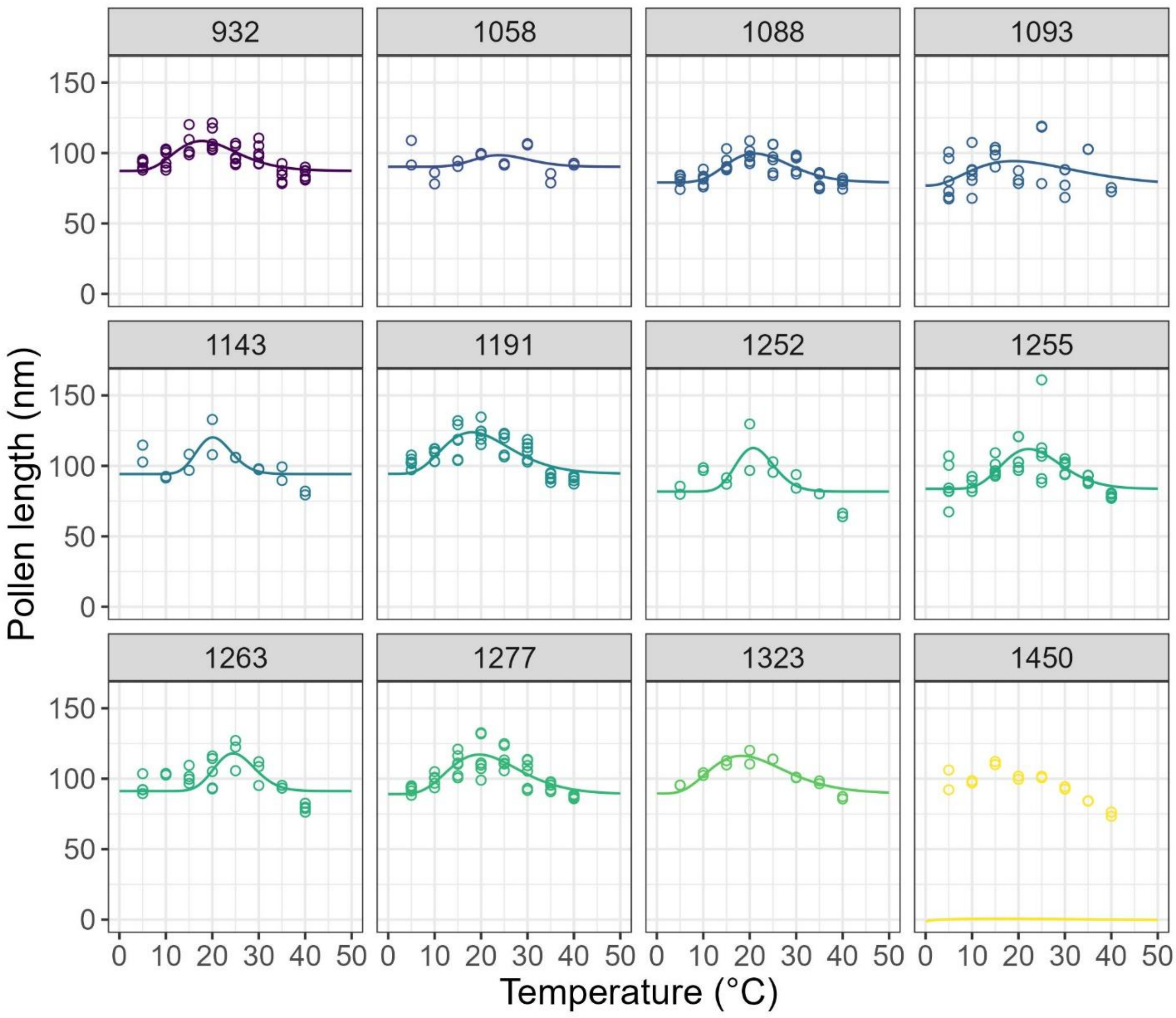


Figure S6. Gamma temperature response curves of ratio-based pollen length (PL) in *Pseudotsuga menziesii* (PSME) from different elevations.

Table S1. Ratio-based pollen germination percentage (GP) and pollen length (PL) optimal temperature of *Pseudotsuga menziesii* (PSME) and germination percentage (GP) and pollen tube length (TL) optimal temperature of *Pinus contorta* (PICO), *Picea engelmannii* (PIEN), and *Pinus ponderosa* (PIPO). Different letters indicate significant differences between species (column-wise comparisons) by using Tukey's honest significance test (HSD). ***$p < 0.001$. Data of PICO, PIEN, and PIPO are from Hsu & Kim (2025).

| | PICO | PIEN | PIPO | PSME | *p*-value |
|---|---|---|---|---|---|
| $T_{opt\_GP}$ (°C) | $23.6 \pm 1.5^{b}$ | $16.4 \pm 1.3^{d}$ | $26.2 \pm 1.7^{a}$ | $22.9 \pm 3.7^{b}$ | <0.001*** |
| $T_{opt\_TL}$ (°C) | $26.5 \pm 1.7^{a}$ | $18.7 \pm 2.0^{c}$ | $26.8 \pm 2.8^{a}$ | $21.1 \pm 3.2^{b}$ | <0.001*** |

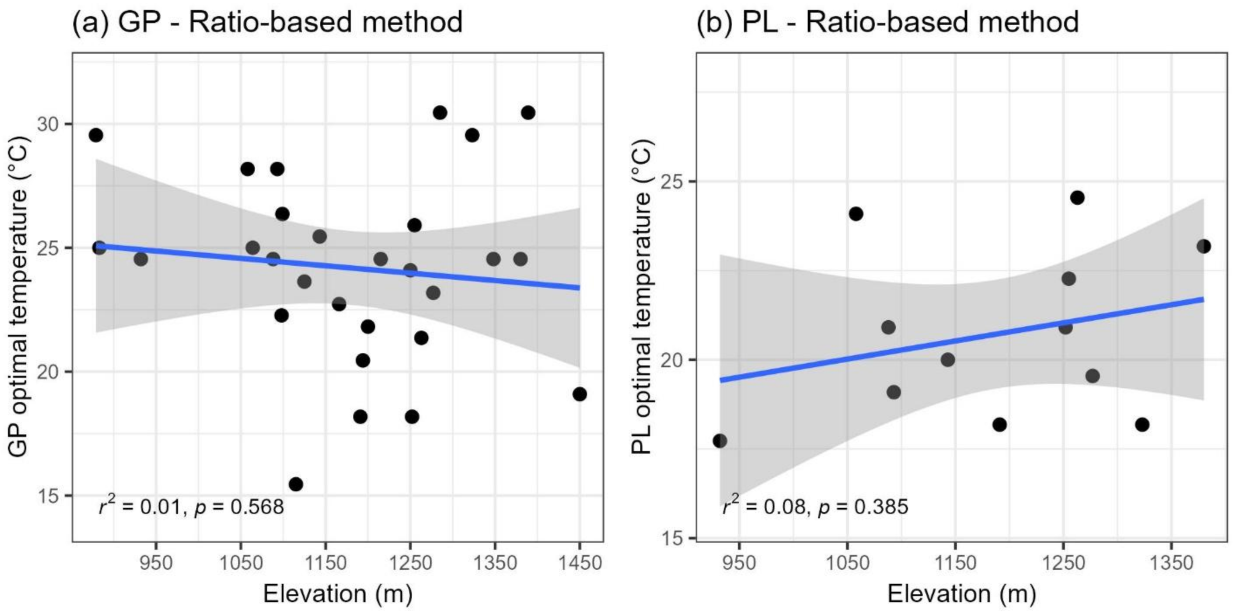


Figure S7. Regression lines of optimal temperatures of ratio-based pollen germination percentage (a; GP) and pollen length (b; PL) in Douglas-fir across the elevational gradients. The shaded areas are 95 % confidence intervals.

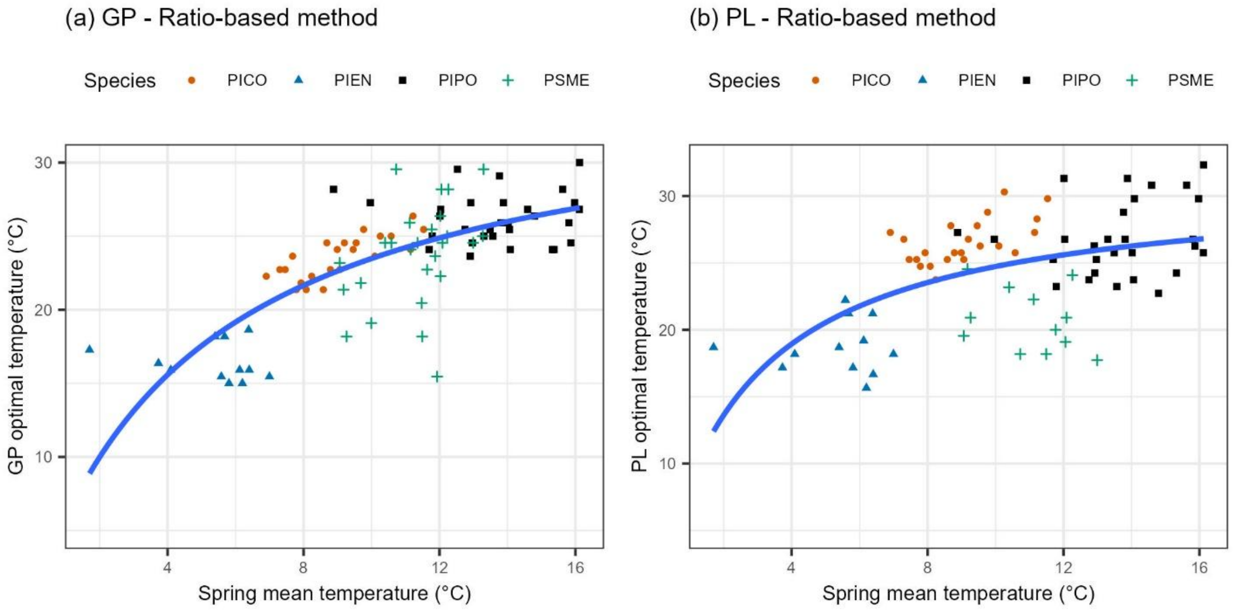


Figure S8. Ratio-based pollen germination percentage (a; GP) and pollen length (b; PL) optimal temperature of *Pseudotsuga menziesii* (PSME) and GP and TL optimal temperature of *Pinus contorta* (PICO), *Picea engelmannii* (PIEN), and *Pinus ponderosa* (PIPO) versus the spring mean temperature across the elevational gradients. The blue lines are Michaelis–Menten equations. Data of PICO, PIEN, and PIPO are from Hsu & Kim (2025).

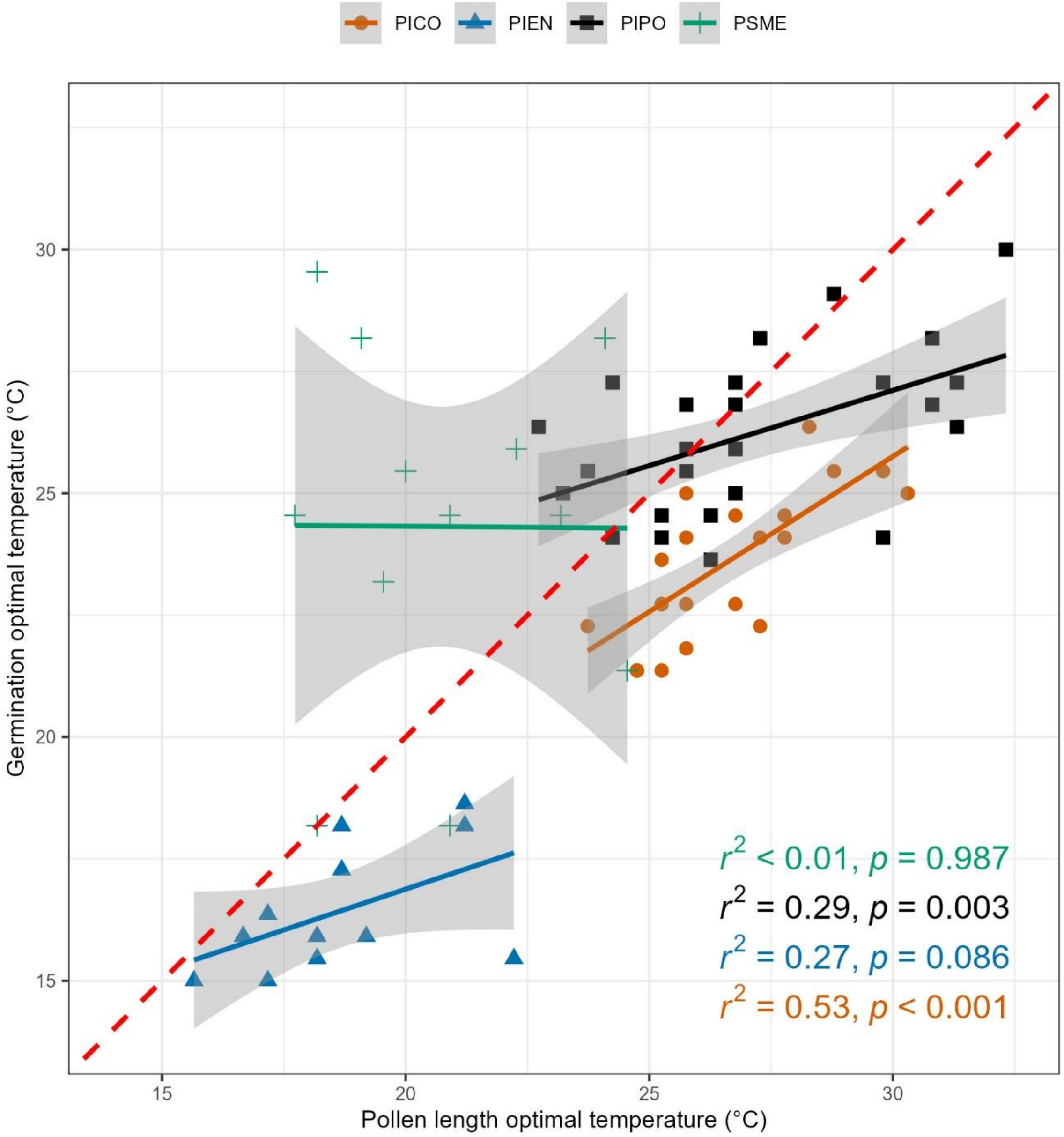


Figure S9. Simple linear regressions between the optimal temperatures of pollen germination percentage (GP) and pollen length (PL) in *Pseudotsuga menziesii* (PSME), and pollen germination percentage and tube length in *Pinus contorta* (PICO), *Picea engelmannii* (PIEN), and *Pinus ponderosa* (PIPO) across the elevational gradients. The red dashed line is the 1:1 temperature ratio. The shaded areas are 95 % confidence intervals. Data of PICO, PIEN, and PIPO are from Hsu & Kim (2025).
